# Polygenic score accuracy in ancient samples: quantifying the effects of allelic turnover

**DOI:** 10.1101/2021.09.21.461259

**Authors:** Maryn O. Carlson, Daniel P. Rice, Jeremy J. Berg, Matthias Steinrücken

## Abstract

Polygenic scores link the genotypes of ancient individuals to their phenotypes, which are often unobservable, offering a tantalizing opportunity to reconstruct complex trait evolution. In practice, however, interpretation of ancient polygenic scores is subject to numerous assumptions. For one, the genome-wide association (GWA) studies from which polygenic scores are derived, can only estimate effect sizes for loci segregating in contemporary populations. Therefore, a GWA study may not correctly identify all loci relevant to trait variation in the ancient population. In addition, the frequencies of trait-associated loci may have changed in the intervening years. Here, we devise a theoretical framework to quantify the effect of this allelic turnover on the statistical properties of polygenic scores as functions of population genetic dynamics, trait architecture, power to detect significant loci, and the age of the ancient sample. We model the allele frequencies of loci underlying trait variation using the Wright-Fisher diffusion, and employ the spectral representation of its transition density to find analytical expressions for several error metrics, including the correlation between an ancient individual’s polygenic score and true phenotype, referred to as polygenic score accuracy. Our theory also applies to a two-population scenario and demonstrates that allelic turnover alone *may* explain a substantial percentage of the reduced accuracy observed in cross-population predictions, akin to those performed in human genetics. Finally, we use simulations to explore the effects of recent directional selection, a bias-inducing process, on the statistics of interest. We find that even in the presence of bias, weak selection induces minimal deviations from our neutral expectations for the decay of polygenic score accuracy. By quantifying the limitations of polygenic scores in an explicit evolutionary context, our work lays the foundation for the development of more sophisticated statistical procedures to analyze both temporally and geographically resolved polygenic scores.

## 1 Introduction

Decay in linkage disequilibrium (LD) between tagging and causal sites, population stratification, variation in allele frequencies within and across populations, and environmental heterogeneity, among other factors, are all thought to negatively impact the prediction accuracy of polygenic scores (see e.g. [1, 2, 3, 4, 5, 6, 7], and more recently in humans, e.g. [8, 9, 10, 11, 12, 13]). Many of these issues likely influence both within- *and* out-of-sample predictions; where out-of-sample may refer to an individual sampled from a distinct time or location relative to that of the GWA study. While empirical [14, 12] and simulation [1, 15, 13] or combined [16] studies have explored particular population genetic scenarios or experimental contexts, we still do not know the extent to which each of these factors compromises prediction accuracy in general.

In this work, we address an issue pertinent to out-of-sample prediction: that causal loci may have different allele frequencies in the GWA study and focal populations. Variants common in the GWA study may be rare in the focal population, and vice versa. We refer to this phenomenon as *allelic turnover*. Allelic turnover implies that effect estimates ported across space and time, or both, may not reflect all of the genetic variation relevant to phenotypic variation in an ancient or geographically distinct population. Allelic turnover further suggests that the statistical properties of ancient polygenic scores depend on when an ancient individual was sampled—a feature not currently accounted for in aDNA analyses. Similarly, statistical properties of geographically disparate polygenic scores depend on the divergence time between GWA study and focal populations. An understanding of allelic turnover in these contexts may ultimately improve statistical analyses of temporally (e.g. [17, 18, 19, 20]) and geographically resolved polygenic scores (e.g. [10, 9]), analyses which are increasingly commonplace.

We aim to quantify the effect of allelic turnover on the polygenic scores of such out-of-sample individuals when they are computed using effect estimates from a contemporary population. We expect that increases in ancient sampling time or divergence time will be associated with declines in polygenic score accuracy due exclusively to allelic turnover. The question is, by how much does accuracy decline? And, can allelic turnover alone explain the reduced accuracy of out-of-sample predictions observed in numerous human (e.g. [16, 15]), animal (e.g. [1, 2, 4]) and plant (e.g. [21, 22]) experiments and simulation studies. The answer is likely to depend on the particular population genetic, trait, and GWA study features of the system under study [3]. We attempt to capture some important aspects of this diversity in our modeling framework.

Here, we consider a standard implementation of the polygenic score *Ŷ* which attributes non-zero effects to a particular set of loci, 𝒮. An individual’s polygenic score is a weighted sum of its genotype, where the weights are the estimated allelic effects. The loci in 𝒮 and their estimated effects are usually identified in large-scale GWA studies, often performed in regional biobanks with sample sizes in the tens to hundreds of thousands of individuals (e.g. the UK Biobank [23], BioBank Japan [24]). Frequently, the set 𝒮 includes loci which are approximately independent and surpass some allele frequency and *p*-value thresholds. Though there are numerous ways to define a polygenic score (e.g. [25, 26]), this implementation is commonly used and proves analytically tractable in our framework.

In contrast to previous quantitative genetic approaches [27, 16], we embed the ancient polygenic score in an explicit population genetic framework. Our theoretical framework allows us to take into account changes in allele frequency as well as the statistical constraint imposed by a finite GWA study sample size. And, distinct from previous approaches to the evolutionary modeling of polygenic scores [28], we track the frequencies of *all loci* that potentially contribute to a trait—not just the loci included in the polygenic score (i.e. loci in 𝒮).

Henceforth, we frame our study exclusively in terms of ancient polygenic scores. However, we formally demonstrate that our theoretical results apply to out-of-space polygenic scores, where the population divergence time multiplied by two is analogous to the ancient sampling time (see Supplementary Text S1.1).

We use several statistics to characterize ancient polygenic score error in distinct population genetic and GWA study scenarios. Each statistic is indexed by the ancient sampling time *τ* : the bias, *bias*(*τ*), mean-squared error, *mse*(*τ*), estimated additive genetic variance, 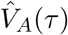, and polygenic score accuracy, *ρ*^2^(*τ*), which approximates the expectation of the squared sample correlation coefficient between polygenic scores and phenotypes in an ancient sample. We derive general forms for these statistics that are agnostic to almost all of our modeling assumptions, and which provide conceptual insights into the effects of allelic turnover. Next, we derive explicit, parameter-dependent expressions for each statistic when the trait is neutrally evolving in a population of constant size subject to recurrent mutation—which for small mutation rates approximates the infinite sites model. We take advantage of the spectral representation of the transition density function of the Wright-Fisher diffusion (*tdf*) to execute these computations [29, 30, 31, 32]. We then find interpretable linear approximations for the initial rate of increase (or decrease) of the metrics under study. These approximations apply for the small ancient sampling times typical of ancient humans remains (e.g. see [18]).

Consistent with our expectations, *mse*(*τ*) increases and the estimated additive genetic variance 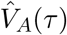 decreases with increasing sampling age *τ*. Despite the fact that *mse*(*τ*) and 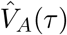 are measuring distinct quantities—and indeed have different functional forms—our linear approximations reveal that, under our assumptions, both statistics initially change at approximately the same rate. This rate is proportional to the product of the mutation rate and the power to detect trait-associated loci in the GWA study, which in turn, is influenced by both study size, the magnitude of the true per-locus effect, and the underlying distribution of the allele frequencies of causal loci.

Moreover, we show that polygenic score accuracy *ρ*^2^(*τ*) is proportional to 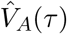, which, as stated, is sensitive to the GWA study and evolutionary parameters. Unlike 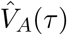, *ρ*^2^(*τ*) depends on the trait heritability *h*^2^, with larger values of *h*^2^ increasing its rate of decay. In contrast, for small mutation rates, relative accuracy, defined as the ratio of *ρ*^2^(*τ*) to accuracy measured in a present-day sample *ρ*^2^(0), is insensitive to *h*^2^, the true per-locus effect size, and the GWA study parameters, as long as the GWA study size *n* exceeds some minimum threshold. We show that this result likely holds for an arbitrary distribution of effects. Importantly, accuracy and relative accuracy decay considerably over the short time spans characteristic of ancient human samples and geographically distinct human populations.

With equal probability of detecting positive versus negative effect alleles, and under neutrality, the bias of the polygenic score is zero for all ancient sampling times. In practice, both of these conditions are likely violated. For example, detection imbalances have been observed in case-control GWA studies [33], and many polygenic traits are likely under some form of selection [34, 35]. Unequal thresholds indeed yield a non-zero *bias*(*τ*) within our framework. But, the magnitude of this bias is small, implying that other perturbations would be necessary to explain any observed, appreciable bias. To relax the neutrality assumption, we simulate recent directional selection. We find that when the selection coefficient is large enough (4*Ns* ≥ 1), selection indeed yields biased polygenic scores. Though this selection-induced bias is several orders of magnitude larger than that induced by asymmetry in the detection thresholds, it is still small relative to the variance explained by segregating genetic variants. Additionally, weak selection only induces small deviations from the neutral theoretical expectations, suggesting that our neutral theory may still accurately capture accuracy declines in the presence of weak directional selection. Altogether, our theoretical results suggest that allelic turnover may make large contributions to out-of-sample reductions in accuracy, even under neutrality.

## 2 Model and metrics

We consider a scenario in which the focal individual is sampled from the same population in which the GWA study was performed, but at a previous point in time *τ*. We specify *τ* in coalescent time units: An ancient sampling time of *τ* corresponds to 2*N* · *τ* generations in the past, with *N* as the population size. When *τ* = 0, the focal individual is an independent sample from the GWA study population.

We summarize the full model in Figure 1 and detail its constituent parts in the proceeding subsections. Briefly, the genotype of the ancient individual is sampled conditional on the population allele frequencies at *τ*. The ancient individual’s phenotype is then sampled conditional on its genotype. Population allele frequencies for all loci that potentially affect the trait evolve until present day, at which point the GWA study is conducted. In particular, the effect sizes included in the polygenic score model are estimated from the genotypes and phenotypes of *n* contemporary individuals. Finally, the ancient polygenic score is computed from the ancient individual’s genotype and the polygenic score model derived from the results of a contemporary GWA study.

**Figure 1:**
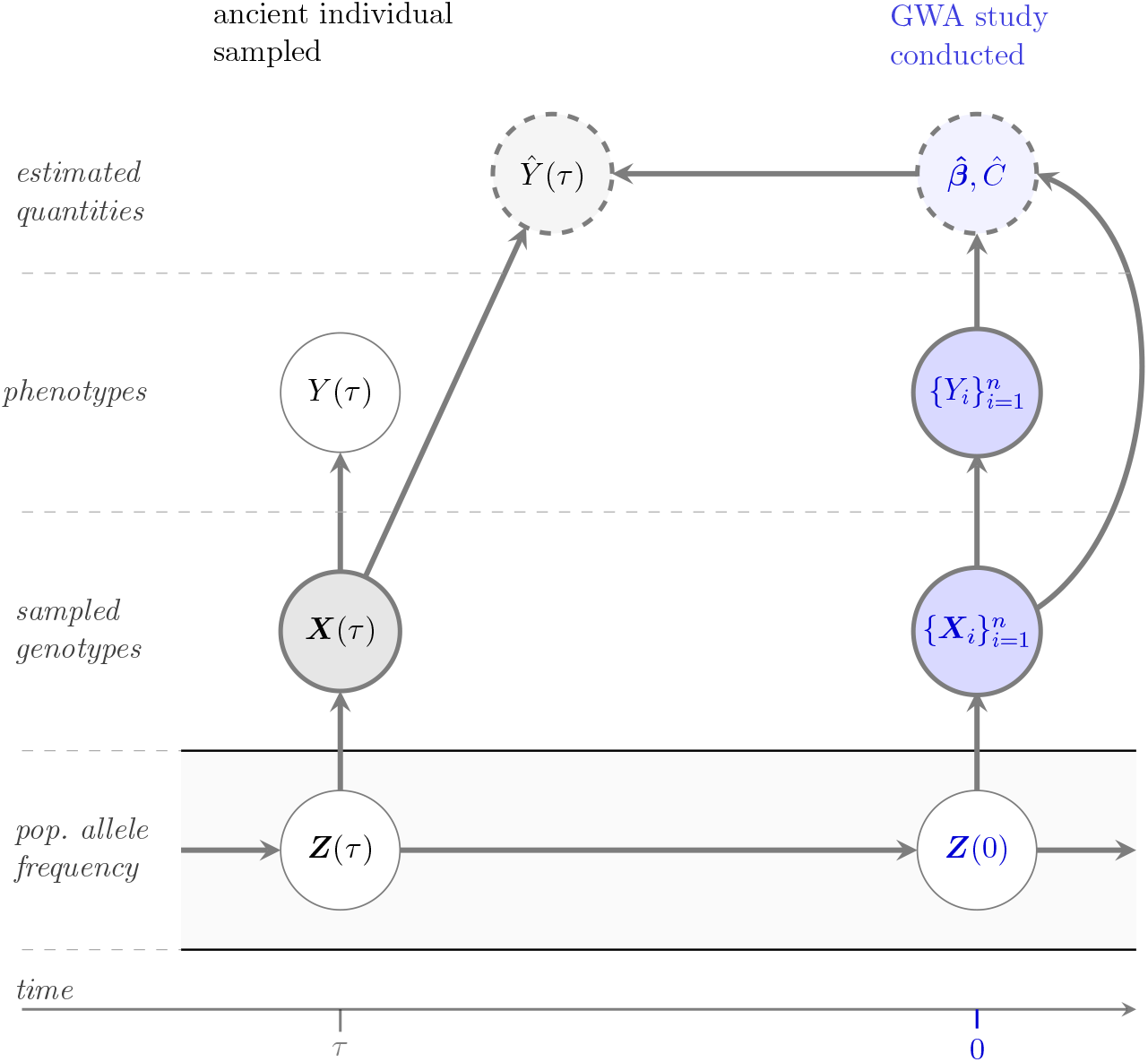
A population genetic model for an ancient polygenic score. A graphical model relating the random variables explicit and implicit in the polygenic score *Ŷ*(*τ*) and phenotype *Y* (*τ*) of an ancient individual sampled *τ* generations in the past. Darkly shaded and thickly bordered nodes are observed quantities. Unshaded and thinly bordered nodes are unobserved. Lightly shaded nodes bordered by dashed lines denote estimated quantities. Edges denote direct dependencies between connected nodes. For example, conditional on the ancient genotype ***X***(*τ*), the polygenic score *Ŷ*(*τ*) is independent of the population allele frequencies ***Z***(*τ*). Quantities in blue are associated with the present day only, and include the population allele frequencies ***Z***(0); the genotypes of the *n* individuals in the GWA study, 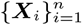 and their phenotypes,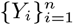; and, the effects and intercept term estimated in the GWA study, 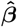 and *Ĉ*, respectively.

### 2.1 Sampling the genotype of a time-indexed individual

We assume that each site is at most bi-allelic, with possible alleles *A*_1_ and *A*_2_. We denote the genotype of an individual sampled at some time *t* (in coalescent units) as *X*_*iℓ*_(*t*), where *i* indexes the individual, and *ℓ* the locus. For the ancient individual(s), *t* = *τ*; for the participants in the GWA study, *t* = 0. For mathematical convenience, we use a symmetric genotype encoding, that is *X*_*iℓ*_(*t*) ∈ {−1, 0, 1}, corresponding to genotypes *A*_1_*A*_1_, *A*_1_*A*_2_, and *A*_2_*A*_2_, respectively. Conditional on the population allele frequency of allele *A*_2_ at *t, Z*_*ℓ*_(*t*), the distribution of *X*_*iℓ*_(*t*) is given by the Hardy-Weinberg sampling probabilities: ℙ{*X*_*iℓ*_(*t*) = −1|*Z*_*ℓ*_(*t*) = *z*} = (1 − *z*)^2^, ℙ{*X*_*iℓ*_(*t*) = 0|*Z*_*ℓ*_(*t*) = *z*} = 2*z*(1 − *z*), and ℙ{*X*_*iℓ*_(*t*) = 1|*Z*_*ℓ*_(*t*) = *z*} = *z*^2^.

### 2.2 Modeling the true phenotype

The genetic basis of a polygenic trait, *Y*, is determined by a set ℒ, consisting of *L* distinct genetic loci (|ℒ| = *L*), each with a true per-locus additive effect *β*_*ℓ*_ ∈ ℝ (for *ℓ* = 1, 2, …, *L*). We further assume that the *L* loci contribute linearly to the trait, such that the true phenotype of the *i*-th individual sampled at *t* is specified by the commonly used additive genetic model [36],

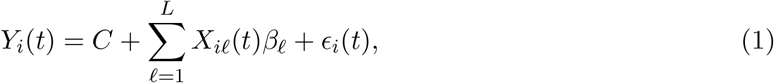

where *C* is a constant; *β*_*ℓ*_ is the true additive effect of locus *ℓ*; and 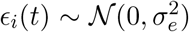 is a normally distributed random variable that incorporates variance in the phenotype due to the environment. The summation in Equation (1) is often referred to as an individual’s *genetic value*. A locus *ℓ* contributes ±*β*_*ℓ*_ to the genetic value (and phenotype) of an individual who is homozygous (at *ℓ*), and zero to that of a heterozygous individual. *C* is thus the phenotype of an hypothetical all heterozygous individual. Without loss of generality, we set *C* = 0. In addition, we assume, without loss of generality, that all *β*_*ℓ*_ ≥ 0 such that locus *ℓ* contributes −*β*_*ℓ*_ to the genetic values of *A*_1_*A*_1_ individuals and +*β*_*ℓ*_ to the genetic values of *A*_2_*A*_2_ individuals.

A fixed locus, *Z*_*ℓ*_(*t*) ∈ {0, 1}, will affect the mean phenotype of the population at *t* by ±*β*_*ℓ*_ but will not contribute to phenotypic variation. We illustrate this fact by conditioning on the allele frequencies of all loci in ℒ at *t*, ***Z***(*t*) ∈ [0, 1]^*L*^. Assuming independence between the environmental and genetic effects, we have,

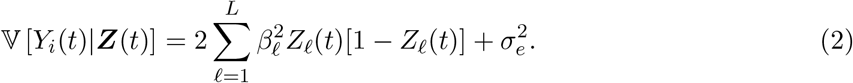

The summation in Equation (2) is the additive genetic variance at *t, V*_*A*_(*t*). For a segregating site, the summand is proportional to *Z*_*ℓ*_(*t*)(1 − *Z*_*ℓ*_(*t*)), with 0 < *Z*_*ℓ*_(*t*)(1 − *Z*_*ℓ*_(*t*)) < 1. For a fixed site, the summand is zero and the site does not contribute to the additive genetic variance *V*_*A*_(*t*). An important feature of our model is that some of the *L* loci may not exhibit genetic variation in the population at a given time. More concretely, the set of loci with non-zero estimated effects on the polygenic score, S, may only be a small subset of ℒ. Thus, we assume that ℒ is a superset of 𝒮.

### 2.3 Constructing a model for the polygenic score

As our aim is to isolate the effects of allelic turnover on the statistical properties of polygenic scores, we make two additional assumptions: (i) the genotyped sites are the causal sites; and (ii) all loci are in linkage equilibrium. Akin to [37], we employ a simple threshold model for the effect estimates. For a GWA study consisting of *n* individuals (and 2*n* chromosomes),

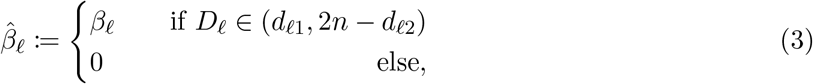

where *D*_*ℓ*_ is the allele count of the trait-increasing allele *A*_2_ at the *ℓ*-th site in the GWA study sample; and *d*_*ℓ*1_ and *d*_*ℓ*2_ are the site-specific detection thresholds. In this simplified model, the true effect is estimated perfectly for all sites with allele counts within the intervals (*d*_*ℓ*1_, 2*n* − *d*_*ℓ*2_) for *ℓ* ∈ ℒ. We allow the two thresholds to differ in order to encompass scenarios in which power is an asymmetric function of the sample allele frequencies, e.g. there is more power to detect low frequency (*D*_*ℓ*_ < *n*) versus high frequency (*D*_*ℓ*_ > *n*) trait-increasing alleles. Such situations may arise with polygenic disease inheritance and imbalanced case and control sample sizes [33]. In most cases, however, we will consider symmetric detection thresholds, with *d*_*ℓ*1_ = *d*_*ℓ*2_ = *d*_*ℓ*_. The threshold *d*_*ℓ*_ depends on on the phenotypic variance, genome-wide significance threshold, true per-locus effect *β*_*ℓ*_, and GWA study size *n*. In Supplementary Text S1.2, we give an explicit form for this dependency for a continuous focal trait and equal detection thresholds, see Equation (S11). Varying *d*_*ℓ*_ while keeping the GWA sample size fixed is equivalent to varying the true per-locus effect *β*_*ℓ*_. Varying the GWA study size *n* while keeping *β*_*ℓ*_ and the other parameters fixed is akin to varying the GWA study’s power to detect loci of a particular effect size. In Section 3, we do both.

The threshold model arises in the large GWA study size *n* limit for the model of 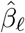 provided in Equation (S5). Namely, as long as *D*_*ℓ*_ is not too small, the variance of 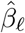 goes to zero as *n* grows. Thus, the threshold model in Equation (3) will necessarily underestimate the true variance of 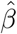 (Supplementary Text S1.4). Still, this model captures the dependency of 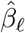 on the GWA study sample size *n* and the true per-locus effect *β*_*ℓ*_, while still facilitating our analytical treatment.

In order to compare the polygenic score with an individual’s true phenotype, we need to account for all sites in the mutational target ℒ, not just those in 𝒮, the set of sites with non-zero effect estimates in the polygenic score. As 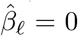 for any site in ℒ but not 𝒮, we express the polygenic score as a function of all loci in ℒ. The ancient polygenic score of individual *i* sampled *τ* generations in the past is then given by,

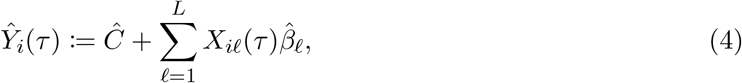

where *Ĉ* is the average phenotype of the GWA sample after subtracting the estimated genetic effects at all loci,

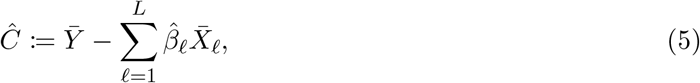

with 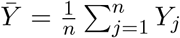 and 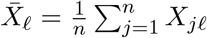 as the mean phenotype and genotype at locus *ℓ* in the GWA study sample, respectively. Here, and in the remainder of our study, we omit time-indexing for random variables associated with the GWA study at *t* = 0. By design, the estimated intercept *Ĉ* absorbs the effects of all loci which were not detected as significant in the GWA study, i.e. those sites for which 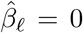. Its presence in the polygenic score of Equation (4) is necessitated by the fact that, to facilitate our analytical treatment, we did not center nor scale the genotypes and phenotypes in the GWA study. Importantly, all of our results are independent of this choice (Supplementary Text S1.5). Henceforth, unless otherwise noted, we refer to Equation (4) as the *polygenic score* and to the summation in Equation (4) as the *genetic prediction*.

### 2.4 Modeling population genetic dynamics

Population genetic processes govern the correlations between allele frequencies at distinct points in time. We model this correlation using the Wright-Fisher diffusion with recurrent mutation. As we assumed all loci were in linkage equilibrium, their allele frequencies evolve forward in time independently, subject to genetic drift and mutation. At each site, alleles mutate from *A*_1_ → *A*_2_ with rate *µ*, and from *A*_2_ → *A*_1_ with rate *ν*. While our results readily generalize to arbitrary *µ* and *ν*, we restrict ourselves to equal mutation rates, *µ* = *ν*.

We further assume that the population is at equilibrium. In this setting, the marginal allele frequencies are beta-distributed, with shape and scale parameters specified by the population-scaled mutation rate; we denote the latter quantity by *a*, with *a* = 4*Nµ* = 4*Nν*.

The relative magnitudes of mutation and genetic drift determine which force dominates an allele frequency trajectory. For example, as *a* approaches 0, the effects of mutation on the frequencies of segregating mutations become negligible and genetic drift dominates. In this low mutation regime (*a* ≪ 1, or equivalently 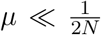), the recurrent mutation model approximates the infinite sites model, while still retaining the features that make it attractive for our analytical treatment. In particular, the stationary allele frequency distribution is a well-defined probability distribution under the recurrent mutation model, but not under the infinite sites model. We concern ourselves almost exclusively with the low mutation regime.

### 2.5 Quantifying *out-of-sample* prediction errors

To quantify how well the polygenic score approximates the true phenotype of an individual randomly sampled from the population *τ* generations ago, we use several statistics:

#### Bias

We define the bias as the expectation of the difference between the polygenic score and true phenotype,

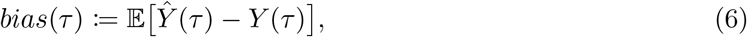

where, here and elsewhere, we omit the subscript when there is only one sample. The expectation in Equation (6) is with respect to the entire random process, encompassing the underlying population genetic dynamics, estimation of the per-locus effects in the GWA study, and computation of the ancient polygenic score (illustrated in Figure 1).

#### Mean-squared error (*mse*)

We define the *mse* as the expectation of the squared prediction error,

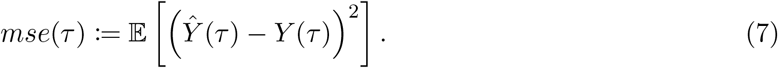

As in Equation (6), the expectation in Equation (7) is with respect to all sources of randomness in the model. The *mse* captures variance in the predictor that exceeds the square of the bias.

#### Expected estimated additive genetic variance 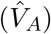

The estimated additive genetic variance is an estimate of the amount of phenotypic variance in the ancient population explained by additive genetic effects alone. We use 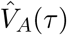 to represent the expectation of this quantity,

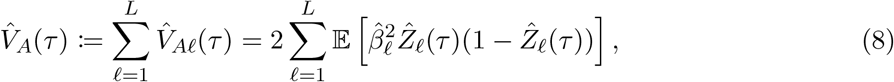

where 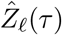 is an estimate of the ancient population allele frequency computed from a sample of *n*_a_ individuals sampled at *τ*. The expected true additive genetic variance, 𝔼[*V*_*A*_], can be found by taking the expectation of the summation in Equation (2).

#### Polygenic score accuracy (*ρ*^2^)

Practitioners often compute the sample correlation coefficient *r*^2^ to measure the accuracy of a predictor in a sample. Here, our sample is *n*_*a*_ ancient individuals sampled from time *τ*, thus,

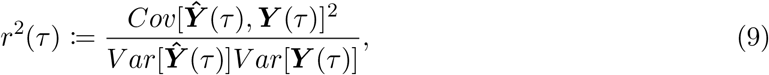

where *Cov*[·, ·] and *V ar*[·] are the sample covariance and variance operators, respectively, and 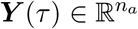 are the *n*_*a*_-dimensional vectors of polygenic scores and phenotypes of the ancient individuals, respectively. Ideally, we would compute the expectation of this quantity—but, this is challenging within our framework due to the common difficulty of computing an expectation of a ratio of random variables. Thus, we approximate the expectation of *r*^2^(*τ*) as the ratio of expectations,

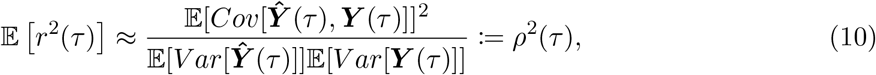

where, as above, the covariance and variances are taken with respect to the sample of *n*_*a*_ ancient individuals, while the expectation is over all sources of randomness in Figure 1 (see Supplementary Text S2.4 for more details). We present simulations in Section 3.4 showing that *ρ*^2^(*τ*) is a good approximation for the expectation of *r*^2^(*τ*) in the parameter regimes of interest.

## 3 Analytical Results

By how much does the prediction accuracy of a polygenic score decrease as the time between sampling the ancient individual and conducting the GWA study increases? To answer this question, we consider a trait potentially influenced by *L* genetic loci, each with true effect *β*_*ℓ*_ ≥ 0, *ℓ* = 1, …, *L*. The forward evolution of sites underlying this trait is modulated by a per site, per generation mutation rate, *µ*, and a population scaled rate of *a* = 4*Nµ*. The diploid population of size 2*N* chromosomes is assumed to be at equilibrium. The parameters dictating the GWA study are the sample size *n* and the detection thresholds specified by ***d***_1_, ***d***_2_ ∈ {1, …, *n*}^*L*^. The metrics are indexed by the ancient sampling time *τ* in coalescent time-units. An ancient sampling time of *τ* corresponds to 2*N* · *τ* generations in the past. We omit the time index for variables associated with the GWA study, which occurs at present day (*t* = 0).

Each subsection is structured as follows: We first derive a general expression for the statistic that does not depend on how we model the population genetic dynamics nor the GWA study. Second, we derive an analytical expression for the statistic under the population genetic assumptions (Section 2.4) and the GWA study threshold model (Section 2.3).

### 3.1 Bias

We can rewrite the sampling time-dependent bias defined in Equation (6) as,

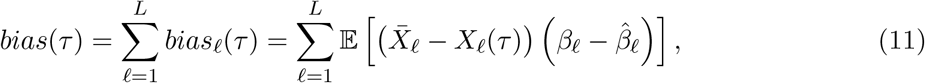

where *bias*_*ℓ*_(*τ*) is the contribution of locus *ℓ* to *bias*(*τ*). From Equation (11), we see that *bias*_*ℓ*_(*τ*) ≈ 0 when either or both of 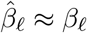 and 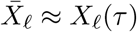 are true. Thus, *bias*_*ℓ*_(*τ*) is minimal when (i) effect estimates are accurate, and (ii) the allele frequencies have not changed substantially in the interval [*τ*, 0].

Under the assumption of equal mutation rates and detection thresholds (*d*_*ℓ*1_ = *d*_*ℓ*2_), *bias*_*ℓ*_(*τ*) = 0 for *τ* ≥ 0 for a reason distinct from those stated above (see Supplementary Text S2.1). Trait-increasing alleles at high frequencies (*D*_*ℓ*_ > *n*) and low frequencies (*D*_*ℓ*_ < *n*) are detected as significant 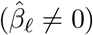 with equal probability. An equivalent assumption is that power is not affected by whether the most prevalent allele is trait-increasing or decreasing. Subsequent evolution of the allele frequencies preserves this symmetry and *bias*(*τ*) remains equal to zero for all *τ*.

However, if we introduce asymmetry in the detection thresholds (*d*_*ℓ*1_ ≠ *d*_*ℓ*2_), *bias*(*τ*) is non-zero for all *τ* (Supplementary Text S2.1). Using the spectral representation of the transition density of the Wright-Fisher diffusion (*tdf*), we derive the per-locus contribution to the bias, *bias*_*ℓ*_(*τ*) (Supplementary Text S2.1). For a small population-scaled mutation rate *a* and a large GWA study size *n*, we approximate this expression, Equation (S38), as,

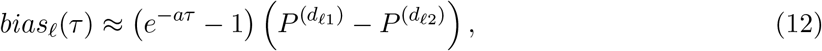

where,

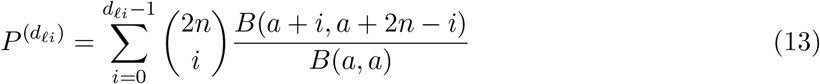

is the probability that the allele count of site *ℓ* is less than *d*_*ℓi*_, i.e. *D*_*ℓ*_ < *d*_*ℓi*_ for *i* = 1, 2; and, *B*(·, ·) is the beta function. Thus, the magnitude of *bias*_*ℓ*_(*τ*) is approximately proportional to the difference in the probability of detecting high (*D*_*ℓ*_ > *n*) versus low (*D*_*ℓ*_ < *n*) frequency alleles, and increases exponentially with *τ*. With a large GWA study size *n* and a small mutation rate *a*, this difference is small relative to the square root of the additive genetic variance—the ratio of these two quantities is smaller than 𝒪(*a*) (Figure S2a). This is due to the fact that when the mutation rate is small, most alleles are close to fixation or fixed. The stationary population allele frequency density *κ*(*z*) ∝ *z*^*a*−1^(1 − *z*)^*a*−1^ behaves like *z*^−1^(1 − *z*)^−1^ for small *a*. Varying *d*_*ℓi*_ then has relatively little impact on 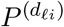, constraining the difference between the one-sided detection probabilities (Figure S2).

### 3.2 Mean-squared error

The sampling time-dependent mean-squared error *mse*(*τ*) can be expressed as,

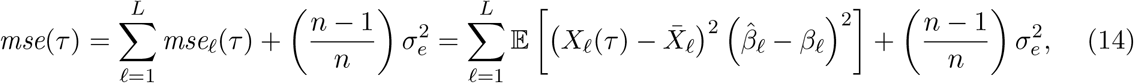

where 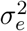 is the variance in the phenotype due to the environment (Supplementary Text S2.2). Note the similarity of the left term in Equation (14) to the form of *bias*(*τ*) given in Equation (11)—similar heuristics apply. Under the threshold model specified in Equation (3), sites at moderate frequencies in the GWA study sample, *D*_*ℓ*_ ∈ [*d*_*ℓ*_, 2*n* − *d*_*ℓ*_], will not contribute to *mse*(*τ*) since 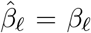. Only sites with frequencies outside this interval (including sites invariant in the GWA study sample) will contribute, and their contributions will be proportional to the squared difference between *X*_*ℓ*_(*τ*) and 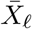. In practice, moderate frequency loci will also contribute to *mse*(*τ*) due to errors in the estimation of the effect estimates and any difference between the ancient genotypes and the average genotypes in the GWA study sample (Supplementary Text S1.4).

We use the spectral representation of the *tdf* (Supplementary Text S1.6) to derive an analytical expression for *mse*_*ℓ*_(*τ*), the per-locus contribution to the *mse* (Supplementary Text S2.2). From this expression, Equation (S43), we derive a linear approximation for the initial per-locus increase in this statistic, Δ*mse*_*ℓ*_(*τ*). With a symmetric detection threshold (*d*_*ℓ*1_ = *d*_*ℓ*2_ = *d*_*ℓ*_) we have,

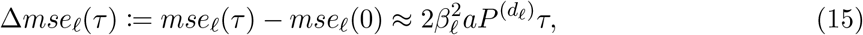

where *mse*_*ℓ*_(0) is the contribution of site *ℓ* to *mse*(*τ*) for *τ* = 0, see Equation (S64); and 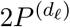, defined in Equation (13), is the probability that the allele count of site *ℓ* is outside the detection interval and 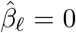. Both *mse*_*ℓ*_(0) and 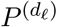 depend on the mutation rate *a*, the GWA study size *n*, and the detection threshold *d*_*ℓ*_.

Δ*mse*_*ℓ*_(*τ*) reflects the time-dependent contributions of sites *not* detected in the GWA study. To see this, we condition on the effect estimate 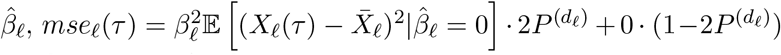. Thus, Equation (15) implies that 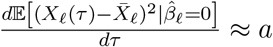 for small *τ*, and consequently, the combined effects of drift and mutation on *mse*_*ℓ*_(*τ*) are captured in the product of the mutation rate and sampling time *aτ*.

In addition, Equation (15) suggests that the rate at which *mse*_*ℓ*_(*τ*) increases will be shared across parameter regimes when 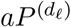 is similar (Figure S3a,d). To illustrate this, we use our analytic formula in Equation (S43) to compute *mse*_*ℓ*_(*τ*) for several low mutation rates, *a* ∈ {10^−4^, 10^−3^, 10^−2^}, and three GWA study sizes, *n* ∈ {10^4^, 10^5^, 10^6^} (Figure 2a). These mutation rates and sample sizes span the range of parameter values appropriate for human data. We depict our results in two ways: (i) we plot the change in *mse*_*ℓ*_(*τ*), and (ii) we plot *mse*_*ℓ*_(*τ*) normalized by the expected additive genetic variance contributed by a single site. At stationarity the expected additive genetic variance is constant and equal to,

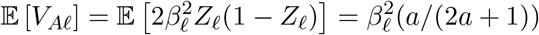

for a scaled-mutation rate *a*. The former plot, Figure 2a, exhibits the functional relationship revealed by Equation (15), while the latter, Figure 2b, approximates the noise-to-signal ratio. In Figure S4, we demonstrate that Equation (15) is a good approximation to *mse*(*τ*) for *τ* ≤ 0.2, particularly when the GWA study size *n* is large.

**Figure 2:**
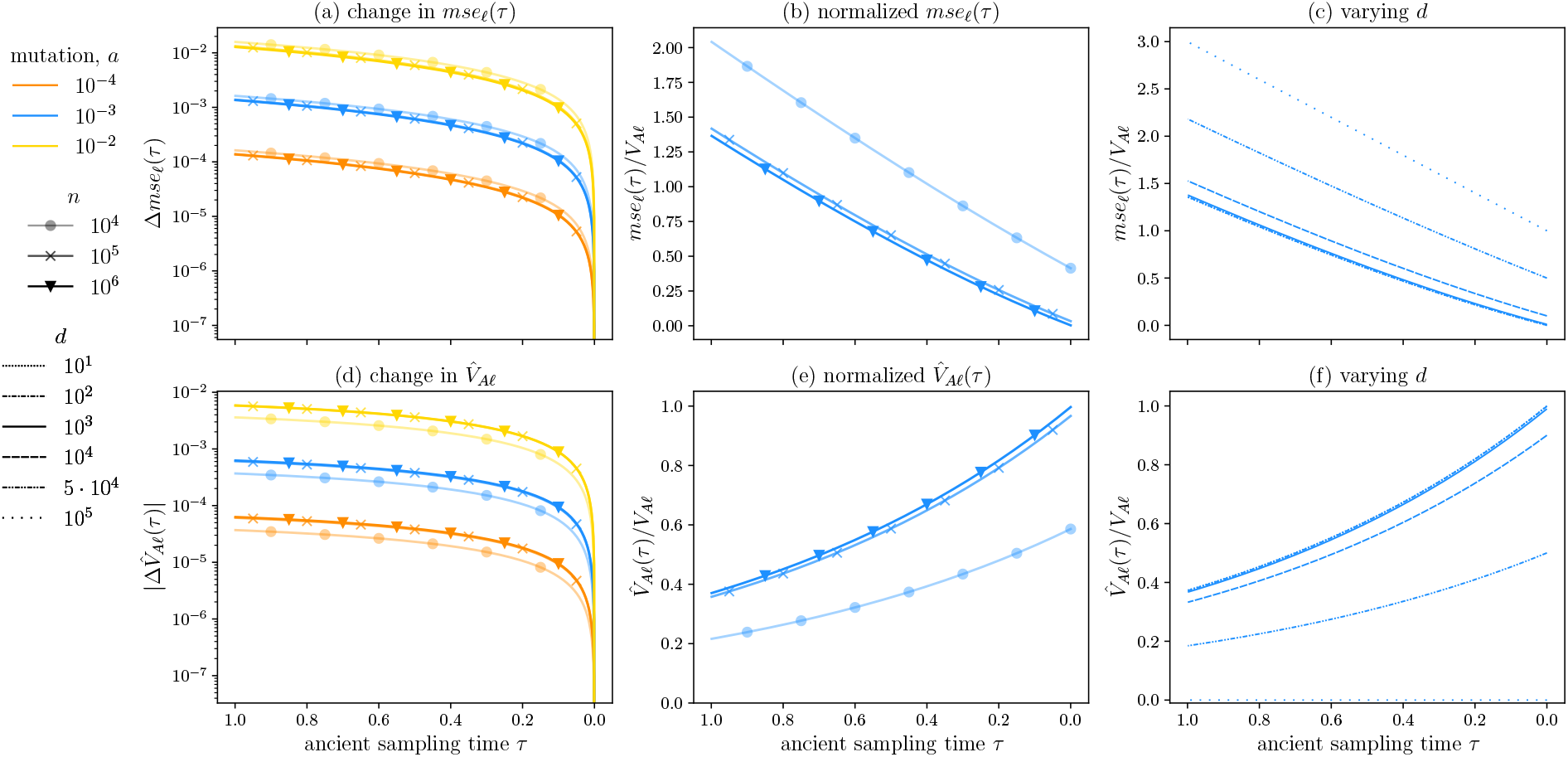
Per locus contributions to the mean-squared error and estimated additive genetic variance across sample sizes, mutation rates, and detection thresholds. In (a), we plot the per-locus increase in *mse*, Δ*mse*_*ℓ*_ (*τ*), normalized by *β*^2^, for three mutation rates *a* = 10^−4^, 10^−3^, 10^−2^ by color, and for the three sample sizes, *n* = 10^4^, 10^5^, 10^6^ by shape, respectively. For a squared effect size of *β*^2^ = 0.01, each sample size, in part, specifies a value of *d*_*ℓ*_, with *d* = 4142, 3340, 3290, or sample allele frequencies of approximately 0.2, 0.02, and 0.002, in order of increasing sample size. In (b-c), we restrict ourselves to *a* = 10^−3^ as the lines for different mutation rates would otherwise largely coincide. In (b), we plot *mse*_*ℓ*_(*τ*) normalized by the expected additive genetic variance at stationarity, 𝔼[*V*_*A*_] = *β*^2^(*a/*2*a* + 1). In (c), we also fix *n* = 10^4^ and vary the detection threshold over several orders of magnitude, *d* ∈ {10, …, 10^5^}, and plot *mse*_*ℓ*_(*τ*) normalized by the expected additive genetic variance. In (d-f), we repeat (a-c), but for the statistic 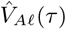, with the following exception: because 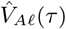 decreases with *τ*, we plot the absolute value of its difference from 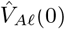 in (a). For all plots the ancient sampling time *τ* ∈ [0, 1], which corresponds to a time span of 2*N* generations.

To find the GWA study size specific detection thresholds used in Figure 2a-b, we solve Equation (S11) for a given effect size *β*, phenotypic variance *V*_*p*_, and significance threshold *α*, while varying the GWA study sample size. For *β*^2^ = 0.01, *V*_*p*_ = 1, and *α* = 10^−8^, the detection thresholds are *d* = 4142, 3340, 3290 in order of increasing sample size, which corresponds to sample allele frequencies of approximately 0.2, 0.02, amd 0.002, respectively. Thus, for a given effect size, larger sample sizes will lead to the detection of alleles at more extreme allele frequencies, while smaller samples will restrict detection to alleles at more intermediate frequencies. Due to non-identifiability, the parameter choices are fairly arbitrary.

We find that for small mutation rates, the cumulative change in the *mse*, Δ*mse*_*ℓ*_(*τ*), is mostly insensitive to differences in the GWA study sample size (Figure 2a,b). The approximation in Equation (15) helps to explain this result. The rate of increase is approximately proportional to 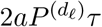. For small mutation rates (*a* ≪ 1) and an arbitrary detection threshold *d*_*ℓ*_, the probability of *not* detecting a locus as significantly associated with the trait is roughly 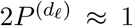 for all sufficiently large *n* (Figure S2b). In this regime, increasing the GWA study sample size only yields small increases in the probability of detecting a locus as significant. Thus, for small mutation rates, the product of this quantity with the mutation rate is 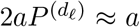, and indeed, we observe a cumulative increase in *mse*_*ℓ*_(*τ*) that is 𝒪(*a*) for *τ* = 1 (Figure 2a). We note that increasing the GWA study sample size does enable detection of loci with smaller effects.

The result in Figure 2a, however, hides the fact that a small absolute increase in *mse*(*τ*) may correspond to a substantial increase in the noise-to-signal ratio. Indeed, for *a* = 10^−3^ (blue lines throughout), *mse*_*ℓ*_(*τ*) ultimately exceeds the expected additive genetic variance E[*V*_*Aℓ*_] for all GWA study sample sizes (Figure 2b). By *τ* = 0.2, a sampling time characteristic of ancient humans, *mse*_*ℓ*_(*τ*) due to allelic turnover is approximately 20% of the additive genetic variance 𝔼[*V*_*Aℓ*_]. For sufficiently large *τ, mse*_*ℓ*_(*τ*) is at least the same order of magnitude as the expected additive genetic variance. In addition, while *mse*_*ℓ*_(*τ*) increases at approximately the same rate irrespective of study size, its initial value *mse*_*ℓ*_(0) is sample size dependent (Figure 2b and see Figure S3b,e for a larger parameter space). Yet, for a given value of *d*_*ℓ*_, reductions in *mse*_*ℓ*_(0) mediated by sample size diminish once *n* is large enough (Figure S3b,e).

Further, Figure 2a obscures the fact that different mutation rates may yield similar noise-to-signal ratios. As discussed, for small *a, mse*_*ℓ*_(*τ*) increases with *τ* at a rate that is 𝒪(*a*). For small *a*, the additive genetic variance is likewise 𝒪(*a*), yielding a relative increase that is mostly insensitive to the mutation rate. Normalized *mse*_*ℓ*_(0) is also similar across small mutation rates (Figure S3b,e), rendering relative *mse*_*ℓ*_(*τ*) mostly insensitive to *a*. We thus omitted the other two mutation rates from Figure 2b.

Lastly, we fix the GWA study sample size at *n* = 10^5^ and vary the detection threshold *d* (Figure 2c). Varying *d* while keeping *n* fixed is analogous to varying the true per-locus effect size *β*, or keeping *β* fixed while varying the significance threshold *α*. The minimum threshold is *d* = 10, whereas *d* = *n* = 10^5^ maximizes *mse*_*ℓ*_(*τ*) since 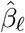 would equal zero for all *ℓ*. Consistent with our analysis above, for small *a*, (i) *mse*_*ℓ*_(0) depends critically on *d*, while (ii) *mse*_*ℓ*_(*τ*)’s approximately linear growth rate is largely insensitive to *d*. Furthermore, by our previous arguments, relative *mse*_*ℓ*_(*τ*) is similar across small mutation rates, and they are also omitted in Figure 2c. For independent and identically distributed (*iid*) loci and 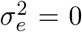, the per-locus *mse*_*ℓ*_(*τ*) values presented in Figure 2b-c are equal to the corresponding trait-wide statistics *mse*(*τ*).

### 3.3 Additive genetic variance

The per-locus contribution to the expected estimated additive genetic variance 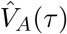 is,

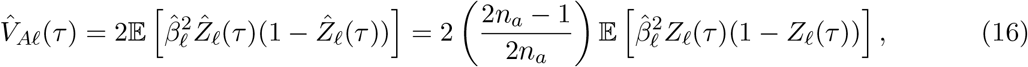

where 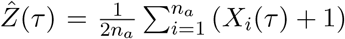 is the estimated allele frequency at *τ*, computed in a sample of size *n*_*a*_ ancient individuals. When 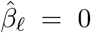 or *Z*_*ℓ*_(*τ*) ∈ {0, 1}, site *ℓ* will not contribute to 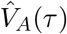. Thus, a site *ℓ* has a non-zero contribution to the estimated additive genetic variance only when it is segregating at both times, the present day and *τ*. This condition is necessary for both 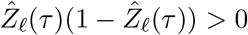 and 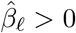 to be true.

As with the two previous statistics, we use the spectral representation of the *tdf* to derive an analytical expression for 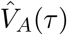 under our population genetic assumptions (Supplementary Text S2.3). The resulting expression, presented in Equation (S47), indicates that the expected additive genetic variance decays exponentially. We then, to first order in the ancient sampling time *τ*, approximate the initial decrease in the per-locus estimated additive genetic variance 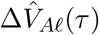,

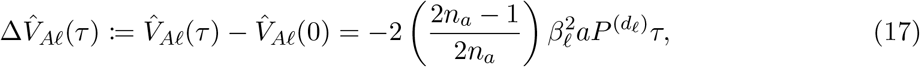

where 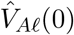 is 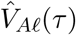 evaluated at *τ* = 0, see Equation (S65); and 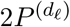, defined in Equation (6), is the probability that 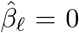. The factor due to finite sampling, 2*n*_*a*_*/*(2*n*_*a*_−1), is ≈ 1 when the ancient sample size *n*_*a*_ is large. Thus, apart from sign, 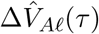 is equal to Δ*mse*_*ℓ*_(*τ*) of Equation (15). Therefore, for small *τ*, 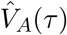 decreases at approximately the same rate as *mse*(*τ*) increases. This result further suggests that for *a* ≪ 1 and a large GWA study size *n*, 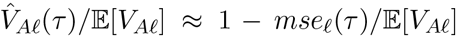 for small *τ* (Figure 2c,f). Although, this relationship trivially breaks down for large *τ* as *mse*_*ℓ*_(*τ*) is not bounded by one.

To compare 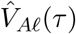 across mutation rates, we mirror our treatment of *mse*_*ℓ*_(*τ*) in the previous section. We plot (i) its increase 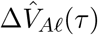 (Figure 2d); (ii) 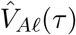 normalized by the expectation of the true additive genetic variance at stationarity, 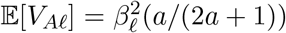 (Figure 2e); and (iii) normalized 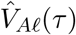, varying the detection threshold for a fixed GWA study sample size (Figure 2f). Akin to *mse*_*ℓ*_(*τ*), normalized 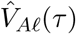 is very similar across small mutation rates. And, while the GWA study size *n* and the detection threshold *d* influence the initial estimated additive genetic variance 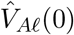, its rate of change is mostly insensitive to the two GWA study parameters.

As 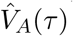 largely recapitulates our results for *mse*(*τ*), with opposing sign, we focus on their differences. Indeed, they have different functional forms and behave differently for modest or large *τ* (see Equation (S47) and Equation (S43), respectively). Conceptually, this discrepancy is not unexpected: In the previous section, we showed that a site only contributes to *mse*(*τ*) if its allele count falls outside the detection interval and 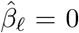. Thus, *mse*(*τ*) increases with *τ* due to alleles shifting from intermediate frequencies in the ancient population to frequencies *outside* of the detection region in the contemporary population. For the expected estimated additive genetic variance 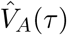, the converse is true: The slope represents the decline in 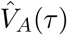 due to alleles changing from frequencies near or at fixation in the ancient population to frequencies *within* the detection interval in the contemporary population. While our results reveal similar functional behavior for these two quantities (with opposing signs) that applies for small *τ*, we caution that statements about 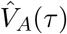 do not immediately translate to statements about *mse*(*τ*), particularly for *τ* ⪆ 0.2.

### 3.4 Polygenic score accuracy

While our framework, in principle, encompasses a trait with varying effect sizes, we will first assume that all sites are *iid* with true effect size *β*. We find in Supplementary Text S2.4 that our approximation to the expectation of the sample correlation coefficient simplifies to,

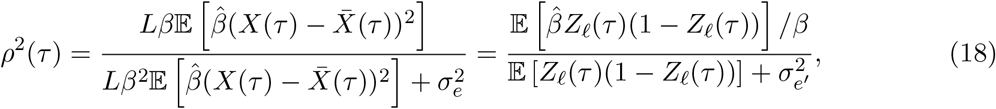

where the compound parameter 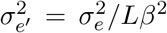 is the environmental variance normalized by the product of the number of loci in the mutational target *L* and the squared per-locus effect size *β*. By comparing Equation (18) with Equation (16), we can see that *ρ*^2^(*τ*) is closely related to the estimated additive genetic variance. Thus, like 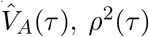 will decrease with *τ* due to loci having changed from frequencies close to zero or one in the ancient population to intermediate frequencies in the contemporary population. However, unlike 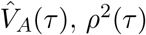 does not depend on the ancient sample size. Therefore, to relate the two statistics, we multiply by the inverse of the ancient sample size dependent factor implicit in 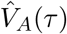,

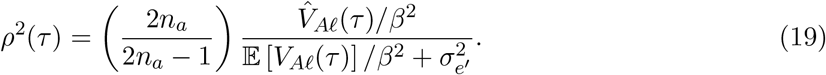

For 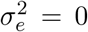, barring the sample size factor, Equation (20) is equal to 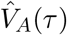 normalized by the expected additive genetic variance. By extension, this quantity approximates the expected sample correlation coefficient *r*^2^(*τ*). A full derivation of Equation (18) from the definition of *r*^2^(*τ*) is provided in Supplementary Text S2.4. By invoking our additional population genetic and GWA study assumptions, we arrive at an approximation for the decrease in polygenic score accuracy,

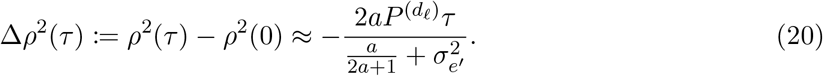

Now, to relate our theory to empirical and simulation studies, one can compute *ρ*^2^(*τ*) for a given narrow-sense heritability *h*^2^ and mutation rate *a* pair. We define *h*^2^ for a trait with a mutational target of *L* loci of equal effects *β*,

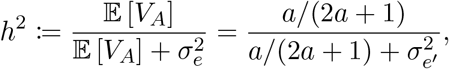

where the equality follows from our population genetic assumptions. Together with *a, h*^2^ fully specifies the compound parameter 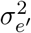 with,

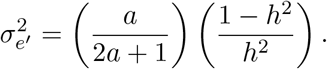

We plot our analytical expressions for both accuracy (Figure 3a) and relative accuracy (Figure 3b), defined as the ratio of *ρ*^2^(*τ*) to *ρ*^2^(0) for *τ* ∈ [1, 0] or 2*N* generations. For humans, this time span corresponds to approximately 500,000 years in the past, encompassing the “Out-of-Africa” migration event, which is estimated to have occurred 50,000-100,000 years ago [38]. We set *h*^2^ = 0.5 and *a* = 10^−3^, and fix the GWA study sample size at *n* = 10^5^. We then compute *ρ*^2^(*τ*), varying the detection threshold over several orders of magnitude (Figure 3a-b). Our results for *ρ*^2^(*τ*) necessarily recapitulate those of 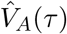: While increasing the detection threshold *d* reduces accuracy substantially (Figure 3a), it does not have a large impact on relative accuracy for *n* = 10^5^ (Figure 3b). Indeed, for small mutation rates, the relative accuracy is insensitive to the mutation rate and threshold, and is well approximated by *e*^−*τ*^, see Equation (S61). Thus, its derivative is also exponential. Absolute accuracy *ρ*^2^(*τ*) likewise decays exponentially, but its derivative is scaled by a quantity that reflects features of the GWA study and the phenotypic variance. For a small mutation rate *a* ≪ 1, its derivative is approximately 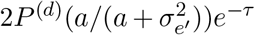, which, in turn, is approximately 2*P* ^(*d*)^*h*^2^*e*^−*τ*^, see Equation (S60). The latter expression suggests that the probability of not detecting a significant association *P* ^(*d*)^ and trait heritability *h*^2^ are the key determinants of prediction accuracy. Importantly, *ρ*^2^(*τ*) declines considerably over the interval *τ* ∈ [1, 0] irrespective of the detection threshold *d*.

**Figure 3:**
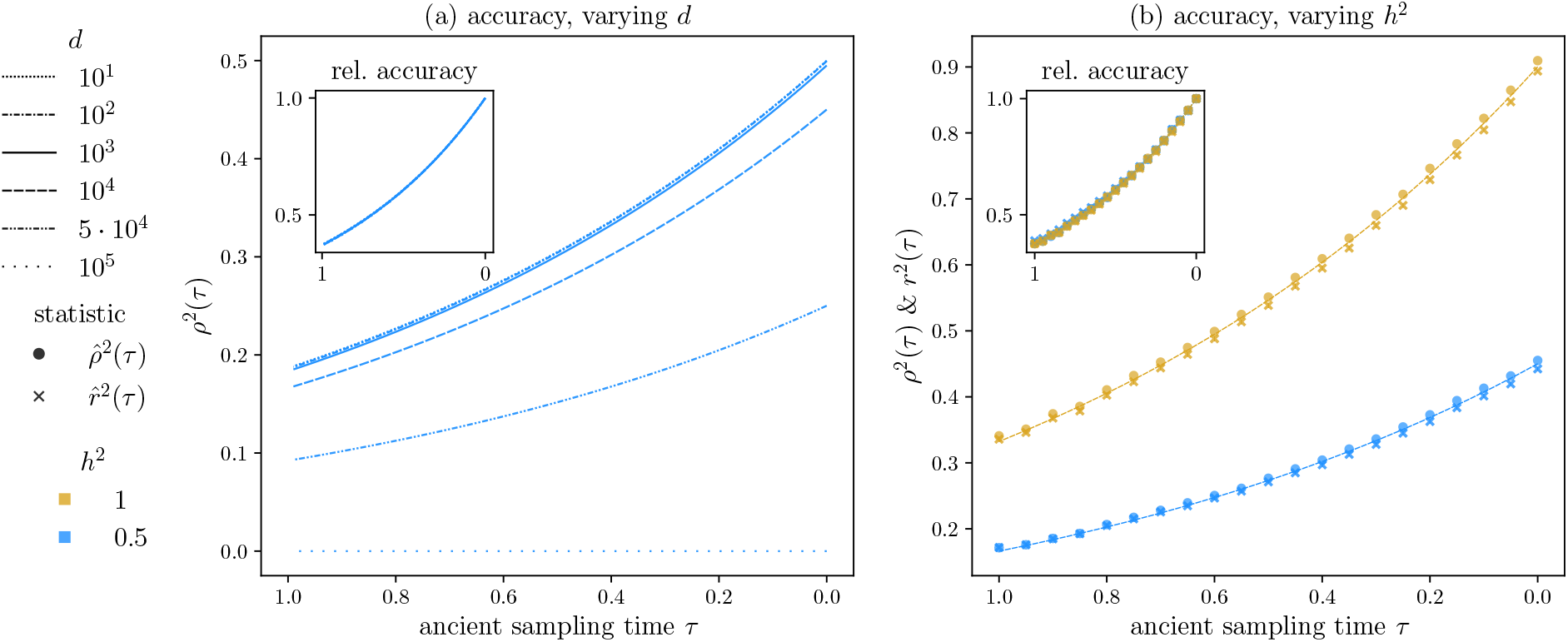
Polygenic score accuracy. We plot our theoretical results for both absolute (a, main) and relative accuracy *ρ*^2^(*τ*) (a, inset) for ancient sampling times *τ* ∈ [0, 1] (or a time span of 2*N* generations) with a mutation rate of *a* = 10^−3^. The GWA study size is shared in all plots and is equal *n* = 10^5^. In (a), we vary the detection threshold over the range of possible values, *d*_*ℓ*_ ∈ {10, … 10^5^}. In (b), we compare our theoretical expectations with simulated estimates of the approximate sample correlation coefficient *ρ*^2^(*τ*) (circles) and the statistic itself *r*^2^(*τ*) (crosses) for a threshold of *d* = 10^4^ (a minimum sample allele frequency of 0.05), and two values of heritability, *h*^2^ = 0.5, 1 (in blue and gold, respectively). The ancient sample size is *n*_*a*_ = 100. In the inset of (b), we normalize the estimates by their initial (estimated) values. Theoretical expressions for *ρ*^2^(*τ*) are also plotted in (b). Each simulated point is the average of *K* = 5000 simulations of *L* = 5000 *iid* loci.

In addition, we glean from Equation (18) that while heritability affects the magnitude of *ρ*^2^(*τ*) through the compound parameter 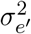, it does not influence the relative accuracy, consistent with previous results [16]. Our simulations suggest that this is also true of the sample correlation coefficient, as our simulated estimates agree extremely well with our theory for *ρ*^2^(*τ*) (Figure 3c-d). We note that this result is contingent on the fact that the environmental variance 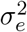 only enters our simple threshold model in the specification of the threshold *d*, see Equation (S11), and does not contribute directly to the variance of the polygenic score (Supplementary Text S2.4). Therefore, we expect this result to hold only for large GWA study sample sizes for which the threshold model is a good approximation to the distribution of 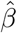. While the finding that relative accuracy is insensitive to the GWA study parameters relies on the assumption that all loci are *iid* and share a causal effect *β*, we provide preliminary theoretical evidence that our results will hold when *β* varies across loci (see Equation (S62) and ensuing comments).

## 4 Simulation results for recent directional selection

We use simulations to explore if and how the statistics under study deviate from their neutral expectations in the presence of recent directional selection. Each copy of the *A*_2_ allele at the *ℓ*-th site confers a fitness advantage of +*s*_*ℓ*_, and so the fitness ratio of the three possible genotypes *A*_1_*A*_1_ : *A*_1_*A*_2_ : *A*_2_*A*_2_ is 1 : 1 + *s*_*ℓ*_ : 1 + 2*s*_*ℓ*_. In our simulations, the population evolves neutrally until the onset of selection at *N* generations (or *τ*_*s*_ = 0.5 in coalescent time units) before present. Thereafter, the population evolves according to discrete Wright-Fisher dynamics with selection.

In the presence of selection, the allele frequency distribution is no longer symmetric; rather, it is skewed toward the beneficial allele. The severity of the skew depends on the selection coefficient and mutation rate, as well as the amount of time that selection has been acting. As we restrict *s*_*ℓ*_ to positive values, designating the *A*_2_ or + allele as beneficial, the allele frequency distribution will be skewed toward one. Because *bias*(*τ*) is proportional to *β*, its sign will be sensitive to this choice. The other statistics will not be affected as long as the detection thresholds are symmetric. Therefore, our results are general up to the sign of *bias*(*τ*).

We conduct simulations over a range of selection coefficients, *σ* = 4*Ns* ∈ {0, 0.1, 1, 10}, for a mutation rate of *a* = 10^−3^. In addition, we plot results for two different detection thresholds, *d* ∈ {10^3^, 10^4^}, in a GWA study sample of size *n* = 10^4^. More details on the simulation procedures are provided in Supplementary Text S1.3.

When *σ* ≥ 1, the polygenic score is biased and positive for *τ* > 0 for both detection thresholds (Figure 4a,b). In other words, with directional selection acting to increase the trait value, *Ŷ*(*τ*) tends to overestimate *Y* (*τ*). The magnitude of *bias*_*ℓ*_(*τ*) depends critically on the strength of selection relative to mutation: We observe a larger bias for *σ* = 10 relative to *σ* = 1, and likewise the bias is larger for *σ* = 1 relative to *σ* = 0.1. In fact, the smaller selection coefficient *σ* = 0.1 is not distinguishable from the neutral expectations. For 0 ≤ *τ* < *τ*_*s*_, *bias*_*ℓ*_(*τ*) increases at an accelerating rate; for *τ* ≥ *τ*_*s*_, *bias*(*τ*) appears constant in this parameter regime.

**Figure 4:**
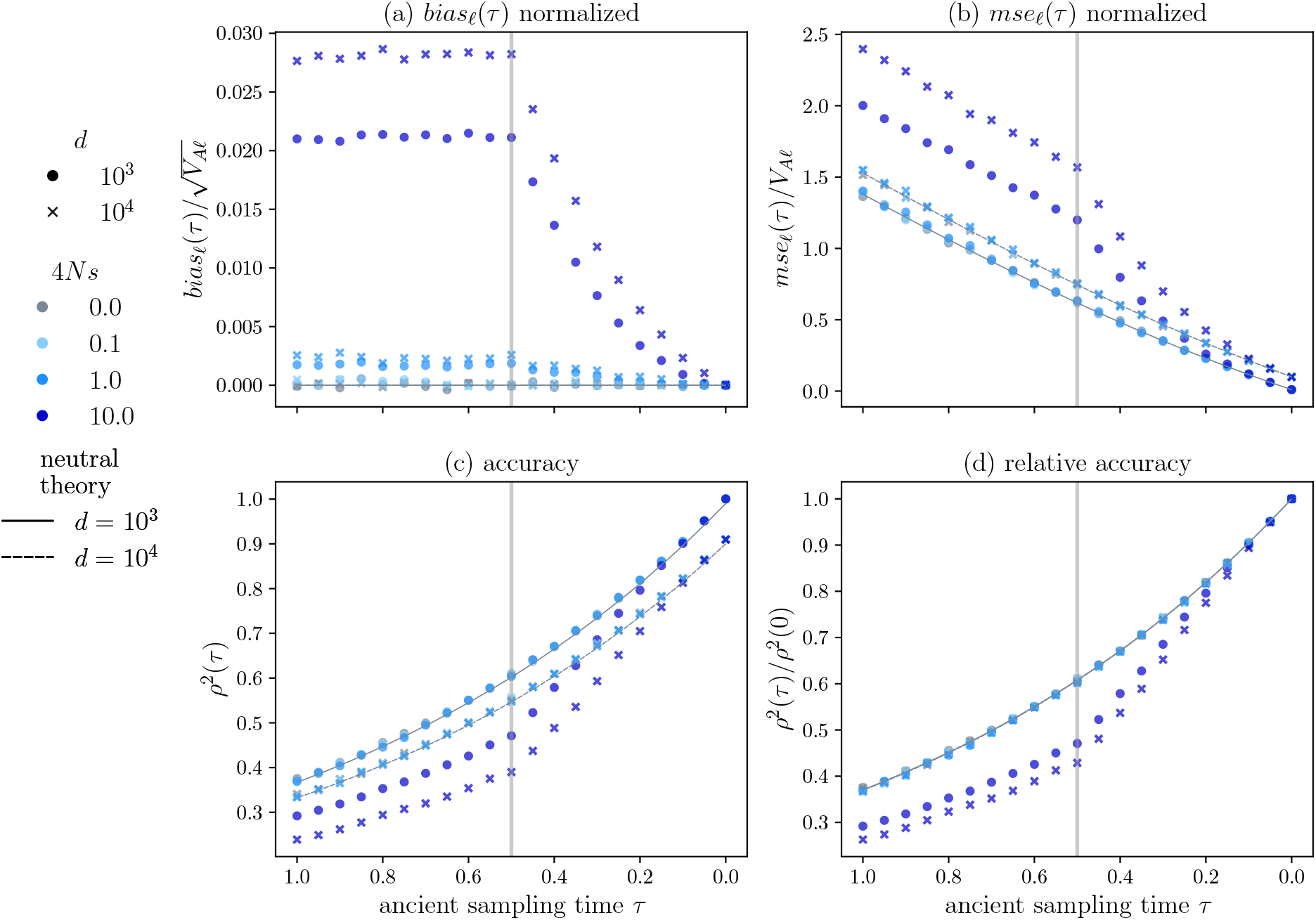
Ancient polygenic scores in the presence of genic selection. We conduct *K* = 5000 simulations, each with a mutational target of *L* = 5000 loci, in a population of size 2*N* = 2 · 10^3^, with a population-scaled mutation rate, *a* = 10^−3^. We consider four selection coefficients, *σ* = 4*Ns* ∈ {0, 0.1, 1, 10} (indicated by color). The GWA study sample size is 2*n* = 2 · 10^5^, with *d* equal to either 10^3^ or 10^4^. In (a-d), we plot the various simulated statistics along with their neutral expectations (solid or dashed black lines). The vertical gray lines indicate the onset of selection at *τ*_*s*_ = 0.5 which corresponds to *N* = 1000 generations. The ancient sample times are *τ* ∈ [1, 0], corresponding to a time span of 2*N* = 2000 generations. We computed, but did not plot, 95% confidence intervals for *bias*(*τ*), *mse*(*τ*), and *r*^2^, as they largely overlapped with the symbols. We note that the oscillations observed in (a) and (b) are not statistically significant.

A higher detection threshold decreases the detection probability. Thus, we expect that the magnitude of *bias*_*ℓ*_(*τ*) will increase with the detection threshold. Indeed, *bias*_*ℓ*_(*τ*) is larger and increases more quickly for the larger detection threshold *d* = 10^4^ compared to *d* = 10^3^ (Figure 4a). Further, our simulations suggest that the detection threshold coupled with the time of the onset of selection govern the magnitude of the bias for *τ* > *τ*_*s*_. For some large *τ, bias*_*ℓ*_(*τ*) will reach an equilibrium value that depends approximately on the asymmetry of the detection thresholds at the present day, which in turn, depends on both the timing and strength of selection (Supplementary Text S2.6).

The underlying allele frequency dynamics provide some insight into these patterns. Before the onset of selection, the allele frequency distribution is stationary and symmetric around 0.5. After the onset of selection, trait-increasing alleles tend to increase in frequency, skewing the distribution toward one. Thus, alleles *not* detected in the GWA study will tend be at higher versus lower frequencies, yielding 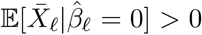 for *σ* > 0. For large *τ*, the frequencies of sites not detected in the GWA study, i.e. with 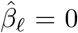, may have substantially changed. Each one of these sites will make a contribution to *bias*(*τ*) that is proportional to 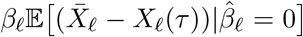 (Equation (11)). Looking backward in time, the shift in the allele frequency distribution ensures that the conditional expectation of *X*_*ℓ*_(*τ*) is smaller than that of 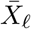, yielding a positive *bias*_*ℓ*_(*τ*) for *τ* > 0. Notably, the magnitude of *bias*_*ℓ*_(*τ*) induced by selection is several orders of magnitude larger than that induced by asymmetry in the detection threshold alone (Figure S2a).

The effects of selection on *mse*_*ℓ*_(*τ*) are qualitatively consistent with those on *bias*_*ℓ*_(*τ*) (Figure 4b). Although, here, the only selection coefficient which induces significant deviations from neutral expectations is *σ* = 10. And, *mse*(*τ*) is larger for *d* = 10^4^ compared to *d* = 10^3^. As with *bias*(*τ*), for 0 ≤ *τ* < *τ*_*s*_, *mse*_*ℓ*_(*τ*) increases at an accelerating rate; before *τ*_*s*_ (*τ* ≥ *τ*_*s*_), *mse*_*ℓ*_(*τ*) appears to increase linearly. Values of *σ* < 10 do not induce noticeable deviations from neutrality for the correlation coefficient *ρ*^2^(*τ*) either. However, strong selection (*σ* = 10) does lead to substantially larger reductions in accuracy relative to our neutral expectations. In addition, for *σ* = 10, relative accuracy is sensitive to the detection threshold, with accuracy decreasing faster for the larger detection threshold.

## 5 Discussion

In this work, we devised a theoretical framework to quantify the effect of allelic turnover on the error and accuracy of out-of-sample polygenic scores. Unlike previous theoretical approaches [16, 27], we averaged over the evolutionary process governing trait evolution, the GWA study from which a polygenic score model is constructed, and the ancient individual’s genotype and phenotype. In doing so, we found explicit expressions for several commonly used metrics that depend on the focal individual’s sampling time, as well as the parameters governing the population genetic dynamics and power to detect trait-associated loci in the GWA study. Mathematical properties of the recurrent mutation model at stationarity enabled us to compute analytical expressions for the metrics of interest under neutrality, and approximations thereof.

Our analytical expressions suggest that allelic turnover alone may be responsible for large reductions in accuracy: For small mutation rates, *ρ*^2^(*τ*) (and *r*^2^(*τ*)) decreases substantially within short time-spans (by about 20 percent in 0.2*N* generations, corresponding to approximately 120,000 years in humans), regardless of the detection threshold. Indeed, while the detection threshold influenced the magnitude of the polygenic score accuracy, relative accuracy was insensitive to this parameter. In other words, under neutrality, relative accuracy is insensitive to the magnitude of the per-locus effect and only depends on the underlying allele frequency distribution. Our theory suggests that this result will hold for arbitrary distributions of the true effect *β*. Although more work is required to fully substantiate this claim.

Selection, however, induces a dependency between an allele’s effect and its frequency, and may thereby render accuracy sensitive to the detection threshold. Our simulations provide preliminary evidence in support of this claim. For a small mutation rate of *a* = 4*Nµ* = 10^−3^ and a large per-locus selection coefficient *σ* = 4*Ns* = 10, relative accuracy was lower for the larger detection threshold of *d* = 10^4^ compared to *d* = 10^3^. Yet, the difference between detection thresholds was small relative to that induced by selection, and was negligible for smaller selection coefficients. Indeed, smaller selection coefficients (*σ* ≤ 1) did not yield significant deviations from our neutral expectations for the *mse*, accuracy, nor relative accuracy. Therefore, excluding strong selection (*σ* > 1), our neutral expectations for these statistics appear to be good approximations to their true values. Our theoretical results thus prove an apt description of temporally-resolved polygenic scores when polygenic adaptation is achieved by concurrent small frequency changes at numerous small effect loci—a plausible scenario [34, 28]. In addition, the simple patterns revealed by our simulations suggest that it may be possible to derive (approximate) analytic expressions for the given metrics in the presence of strong selection, when loci exhibit selective sweep-like behavior.

It is unclear whether our neutral expectations will hold in the context of more sophisticated polygenic trait modeling. In our simulation study, as in our theoretical work, we focus on dynamics at a single locus. Thus, our results are most relevant to scenarios in which single locus dynamics can be decoupled from the evolution of the mean phenotype and the genetic background [39]. Namely, the effect of an individual locus must be small relative to the mean phenotype [39, 37]. Future work will assess polygenic score accuracy under more sophisticated models of polygenic adaptation (e.g. [37, 40]).

Of the two bias-inducing processes explored, detection threshold asymmetry and directional selection, the latter induced much larger deviations from our neutral expectation for the bias, i.e. under neutrality *bias*(*τ*) = 0 for all ancient sampling times *τ*. In the presence of detection asymmetry, *bias*(*τ*) is approximately proportional to the difference between the one-sided detection probabilities. This difference is constrained by the shape of the stationary allele frequency density. Under neutrality, and for small mutation rates, most alleles are at very low frequencies or fixed, such that changing the detection threshold minimally influences the one-sided detection probabilities. Selection, however, perturbs the underlying allele frequency density. At equilibrium, this density is proportional to *e*^*σz*^*z*^−1^(1 − *z*)^−1^ for small *a*, where *σ* = 4*Ns*. Depending on *σ*, the one-sided detection probabilities may differ markedly, yielding larger values of *bias*(*τ*). We thus suspect that detection asymmetry has the potential to further exacerbate any bias induced by selection. These results are interesting in light of those of Chan et al. 2014 [33], who demonstrated that polygenic disease inheritance under the liability threshold model induced differences in the power to detect protective versus susceptible alleles. This effect was further increased by imbalances in the case and control sample sizes in the GWA study.

Our results clarify relationships between various commonly used metrics of prediction error and accuracy. For example, we demonstrated an approximate functional relationship between the mean-squared error *mse*(*τ*) and the expected additive genetic variance 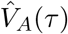 that applies for small ancient sampling times and mutation rates. This shared initial rate emerged despite fundamental differences between these statistics: *mse*(*τ*) measures error due to variants near or at fixation in the contemporary sample, which were segregating at intermediate frequencies in the ancient sample. In contrast, 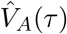 measures error due to variants segregating in the contemporary sample, which were near or at fixation in the ancient sample. This conceptual result does not rely on any of our population genetic or GWA modeling assumptions, and perhaps could be exploited to learn about the genetic architecture of quantitative traits from multi-population data. In addition, we showed formally that polygenic score accuracy *ρ*^2^(*τ*), an approximation to the expectation of the sample correlation coefficient *r*^2^(*τ*), is proportional to the ratio of 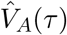 to the total phenotypic variance. We believe that these relations, and their evolutionary and GWA study dependent forms, may facilitate the development of novel, more principled statistical procedures for the analysis of out-of-sample polygenic scores.

At the same time, the simplifying assumptions underlying our results indicate that significant challenges remain. Indeed, allelic turnover cannot explain all of the reductions in accuracy observed in out-of-sample predictions. For example, to achieve the same accuracy reductions observed in both simulated, e.g. [15] and empirical, e.g. [14, 41, 16], studies of cross-population polygenic scores for contemporary humans, allelic turnover under neutrality would require population divergence times that far exceed their estimated values. Differences in linkage disequilibrium between contemporary human populations may largely explain this discrepancy as most trait-associated loci are likely to be tagging rather than causal sites [16, 12]. As with geographically distinct populations, if LD between the genotyped and causal sites differed in the ancient population, then polygenic score accuracy would suffer [1]. We did not model this effect and assumed that the genotyped site was the causal site. For small ancient sampling or population divergence times, high marker density in the GWA study may mitigate accuracy losses due to LD decay, but, more theoretical work is required to substantiate this claim. While our framework can readily incorporate LD, it is difficult to obtain analytical results when the genotyped marker is *not* the causal site. In lieu of theoretical results, large-scale simulations in simple population genetic scenarios may provide insight into the relative contributions of LD—which depends on the allele frequencies of the tagging and causal sites—and allelic turnover to declines in polygenic score accuracy. In addition, we assumed that per-locus causal effects were shared by the ancient and contemporary samples. Indeed, differences in causal effects across contemporary populations likely contributes to accuracy reductions [8, 12]. We conjecture that fluctuations in the per-locus effects would increase *mse*(*τ*) and decrease accuracy, but not profoundly alter our conclusions. Perhaps, if the fluctuations were asymmetric, e.g. effect sizes tended to increase in time, then *bias*(*τ*) may be non-zero under neutrality. Lastly, technical challenges inherent to the extraction and sequencing of ancient DNA often result in noisy estimates of the ancient genotypes. This additional source of randomness is likely to reduce accuracy and increase *mse*(*τ*), but otherwise should not substantially alter our conclusions.

## Code availability

All of the code developed to produce the figures and simulations in this paper is available in the github repository: https://github.com/marync/ancient_polygenic.

## Acknowledgments

We thank members of the Berg, Novembre, Steinrücken labs, and the Cummings fourth floor for helpful discussions throughout the development of this project. In addition, we thank Jennifer Blanc, Adam Fine, Evan Koch, Zachary Miller, and John Novembre for comments on earlier (or very early) versions of this manuscript. We also give a special thanks to Carlos Marcelo Serván and Micol Tresoldi for numerous insightful discussions over the course of this project.

## References

1. Habier D, Fernando RL, Dekkers JCM. The impact of genetic relationship information on genome-assisted breeding values. Genetics. 2007;177(4):2389–2397. https://doi.org/10.1534/genetics.107.081190.

2. De Roos APW, Hayes BJ, Spelman RJ, Goddard ME. Linkage disequilibrium and persistence of phase in Holstein-Friesian, Jersey and Angus cattle. Genetics. 2008;179(3):1503–1512. https://doi.org/10.1534/genetics.107.084301.

3. Hamblin MT, Buckler ES, Jannink JL. Population genetics of genomics-based crop improvement methods. Trends in Genetics. 2011;27(3):98–106. https://doi.org/10.1016/j.tig.2010.12.003.

4. Erbe M, Hayes BJ, Matukumalli LK, Goswami S, Bowman PJ, Reich CM, et al. Improving accuracy of genomic predictions within and between dairy cattle breeds with imputed highdensity single nucleotide polymorphism panels. Journal of Dairy Science. 2012;95(7):4114–4129. https://doi.org/10.3168/jds.2011-5019.

5. Carlson CS, Matise TC, North KE, Haiman CA, Fesinmeyer MD, Buyske S, et al. Generalization and Dilution of Association Results from European GWAS in Populations of Non-European Ancestry: The PAGE Study. PLoS Biology. 2013;11(9):e1001661. https://doi.org/10.1371/journal.pbio.1001661.

6. Wray NR, Yang J, Hayes BJ, Price AL, Goddard ME, Visscher PM. Pitfalls of predicting complex traits from SNPs. Nature Reviews Genetics. 2013;14(7):507–515. https://doi.org/10.1038/nrg3457.

7. Guo Z, Tucker DM, Basten CJ, Gandhi H, Ersoz E, Guo B, et al. The impact of population structure on genomic prediction in stratified populations. TAG Theoretical and applied genetics Theoretische und angewandte Genetik. 2014;127(3):749–762. https://doi.org/10.1007/s00122-013-2255-x.

8. Galinsky KJ, Reshef YA, Finucane HK, Loh PR, Zaitlen N, Patterson NJ, et al. Estimating cross-population genetic correlations of causal effect sizes. Genetic Epidemiology. 2019;43(2):180–188. https://doi.org/10.1002/gepi.22173.

9. Berg JJ, Harpak A, Sinnott-Armstrong N, Joergensen AM, Mostafavi H, Field Y, et al. Reduced signal for polygenic adaptation of height in UK Biobank. eLife. 2019;8. https://doi.org/10.7554/elife.39725.

10. Sohail M, Maier RM, Ganna A, Bloemendal A, Martin AR, Turchin MC, et al. Polygenic adaptation on height is overestimated due to uncorrected stratification in genome-wide association studies. eLife. 2019;8:1–17. https://doi.org/10.7554/elife.39702.

11. Mostafavi H, Harpak A, Agarwal I, Conley D, Pritchard JK, Przeworski M. Variable prediction accuracy of polygenic scores within an ancestry group. eLife. 2020;9. https://doi.org/10.7554/eLife.48376.

12. Bitarello BD, Mathieson I. Polygenic scores for height in admixed populations. G3: Genes, Genomes, Genetics. 2020;10(11):4027–4036. https://doi.org/10.1534/g3.120.401658.

13. Durvasula A, Lohmueller KE. Negative selection on complex traits limits phenotype prediction accuracy between populations. American Journal of Human Genetics. 2021;108(4):620–631. https://doi.org/10.1016/j.ajhg.2021.02.013.

14. Martin AR, et al. Human Demographic History Impacts Genetic Risk Prediction across Diverse Populations. American Journal of Human Genetics. 2017;100:635–649.

15. Ragsdale AP, Nelson D, Gravel S, Kelleher J. Lessons Learned from Bugs in Models of Human History. American Journal of Human Genetics. 2020;107(4):583–588. https://doi.org/10.1016/j.ajhg.2020.08.017.

16. Wang Y, Guo J, Ni G, Yang J, Visscher PM, Yengo L. Theoretical and empirical quantification of the accuracy of polygenic scores in ancestry divergent populations. Nature Communications. 2020;11(1). https://doi.org/10.1038/s41467-020-17719-y.

17. Swarts K, Gutaker RM, Benz B, Blake M, Bukowski R, Holland J, et al. Genomic estimation of complex traits reveals ancient maize adaptation to temperate North America. Science. 2017;357(6350):512–515. https://doi.org/10.1126/science.aam9425.

18. Cox SL, Ruff CB, Maier RM, Mathieson I. Genetic contributions to variation in human stature in prehistoric Europe. Proceedings of the National Academy of Sciences of the United States of America. 2019;116(43):21484–21492. https://doi.org/10.1073/pnas.1910606116.

19. Colbran LL, Gamazon ER, Zhou D, Evans P, Cox NJ, Capra JA. Inferred divergent gene regulation in archaic hominins reveals potential phenotypic differences. Nature Ecology and Evolution. 2019;3(11):1598–1606. https://doi.org/10.1038/s41559-019-0996-x.

20. Cox SL, Moots H, Stock JT, Shbat A, Bitarello BD, Haak W, et al. Predicting skeletal stature using ancient DNA. bioRxiv. 2021; p. 2021.03.31.437877. https://doi.org/10.1101/2021.03.31.437877.

21. Windhausen VS, Atlin GN, Hickey JM, Crossa J, Jannink JL, Sorrells ME, et al. Effectiveness of genomic prediction of maize hybrid performance in different breeding populations and environments. G3: Genes, Genomes, Genetics. 2012;2(11):1427–1436. https://doi.org/10.1534/g3.112.003699.

22. Lorenz AJ, Smith KP, Jannink JL. Potential and optimization of genomic selection for Fusarium head blight resistance in six-row barley. Crop Science. 2012;52(4):1609–1621. https://doi.org/10.2135/cropsci2011.09.0503.

23. Bycroft C, Freeman C, Petkova D, Band G, Elliott LT, Sharp K, et al. The UK Biobank resource with deep phenotyping and genomic data. Nature. 2018;562(7726):203–209. https://doi.org/10.1038/s41586-018-0579-z.

24. Kanai M, Akiyama M, Takahashi A, Matoba N, Momozawa Y, Ikeda M, et al. Genetic analysis of quantitative traits in the Japanese population links cell types to complex human diseases. Nature Genetics. 2018;50(3):390–400. https://doi.org/10.1038/s41588-018-0047-6.

25. Meuwissen TH, Hayes BJ, Goddard ME. Prediction of total genetic value using genome-wide dense marker maps. Genetics. 2001;157(4):1819–1829. https://doi.org/10.1093/genetics/157.4.1819.

26. de los Campos G, Hickey JM, Pong-Wong R, Daetwyler HD, Calus MPL. Whole-genome regression and prediction methods applied to plant and animal breeding. Genetics. 2013;193(2):327–345. https://doi.org/10.1534/genetics.112.143313.

27. Daetwyler HD, Villanueva B, Woolliams JA. Accuracy of predicting the genetic risk of disease using a genome-wide approach. PLoS ONE. 2008;3(10). https://doi.org/10.1371/journal.pone.0003395.

28. Berg JJ, Coop G. A Population Genetic Signal of Polygenic Adaptation. PLoS Genetics. 2014;10(8):1004412. https://doi.org/10.1371/journal.pgen.1004412.

29. Ewens WJ. Mathematical Population Genetics I: Theoretical Introduction. New York: Springer-Verlag; 2004.

30. Durrett R. Probability Models for DNA Sequence Evolution. 2nd ed. New York: Springer-Verlag; 2008.

31. Griffiths RC, Spano D. Diffusion processes and coalescent trees. arXiv. 2010. http://arxiv.org/abs/1003.4650.

32. Song YS, Steinrücken M. A simple method for finding explicit analytic transition densities of diffusion processes with general diploid selection. Genetics. 2012;190(3):1117–1129. https://doi.org/10.1534/genetics.111.136929.

33. Chan Y, Lim ET, Sandholm N, Wang SR, McKnight AJ, Ripke S, et al. An excess of risk-increasing low-frequency variants can be a signal of polygenic inheritance in complex diseases. American Journal of Human Genetics. 2014;94(3):437–452. https://doi.org/10.1016/j.ajhg.2014.02.006.

34. Pritchard JK, Pickrell JK, Coop G. The Genetics of Human Adaptation: Hard Sweeps, Soft Sweeps, and Polygenic Adaptation. Current Biology. 2010;20(4):208–215. https://doi.org/10.1016/j.cub.2009.11.055.

35. Boyle EA, Li YI, Pritchard JK. An Expanded View of Complex Traits: From Polygenic to Omnigenic. Cell. 2017;169(7):1177–1186. https://doi.org/10.1016/j.cell.2017.05.038.

36. Lynch M, Walsh B. Genetics and Analysis of Quantitative Traits. 1st ed. Sinauer Associates; 1998.

37. Simons YB, Bullaughey K, Hudson RR, Sella G. A population genetic interpretation of GWAS findings for human quantitative traits. PLoS Biology. 2018;16. https://doi.org/10.1371/journal.pbio.2002985.

38. Jouganous J, Long W, Ragsdale AP, Gravel S. Inferring the joint demographic history of multiple populations: Beyond the diffusion approximation. Genetics. 2017;206(3):1549–1567. https://doi.org/10.1534/genetics.117.200493.

39. Chevin LM, Hospital F. Selective sweep at a quantitative trait locus in the presence of background genetic variation. Genetics. 2008;180(3):1645–1660. https://doi.org/10.1534/genetics.108.093351.

40. Hayward LK, Sella G. Polygenic adaptation after a sudden change in environment. bioRχiv. 2019. https://www.biorxiv.org/content/10.1101/792952v2.

41. Duncan L, Shen H, Gelaye B, Meijsen J, Ressler K, Feldman M, et al. Analysis of polygenic risk score usage and performance in diverse human populations. Nature Communications. 2019;10(1). https://doi.org/10.1038/s41467-019-11112-0.

