## Supplementary Text (S1 & S2) for "Polygenic score accuracy in ancient samples: quantifying the effects of allelic turnover"

### Supplementary Information for "Polygenic score accuracy in ancient samples: quantifying the effects of allelic turnover"

Maryn O. Carlson[1*], Daniel P. Rice[2], Jeremy J. Berg[1,2], and Matthias Steinrücken[1,2,3*]

[1]Committee on Genetics, Genomics, & Systems Biology, University of Chicago, Chicago, IL, USA
[2]Department of Human Genetics, University of Chicago, Chicago, IL, USA
[3]Department of Ecology & Evolution, University of Chicago, Chicago, IL, USA

September 21, 2021

### S1 Extended model and methods

Here, we provide additional modeling details and support for claims made in the main text.

#### S1.1 The joint allele frequency density in the population split scenario

In the main text, we claim that under our assumptions—namely, neutrality, constant population size, and stationarity—the modeling framework readily encompasses a simple population split scenario in which two populations diverged some $\tau_{\text{split}}$ generations ago and the ancient individual was sampled at $\tau$ (Figure S1b). Specifically, the split scenario is analogous to the single population scenario (Figure S1a) in which the ancient individual is sampled at $2\tau_{\text{split}} - \tau$. We derive the form of the joint allele frequency density for the demographic split scenario (Figure S1b) as proof.

(a) A single population.

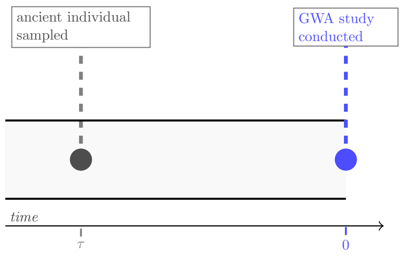

(b) A population split.

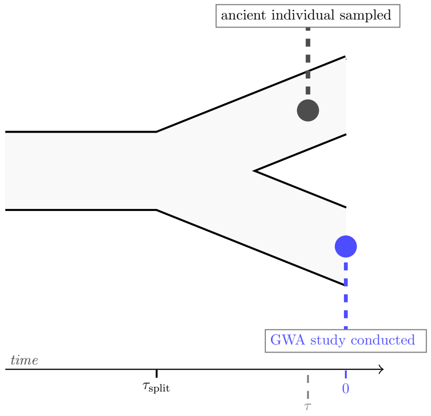

Figure S1: The two demographic and sampling scenarios of interest.

For the scenario in Figure S1a, the joint allele frequency density at $\tau$ and the present is $f(z_\tau, z_0) = p(z_\tau, z_0; \tau)\kappa(z_\tau)$, where $\kappa(\cdot)$ is the stationary density,

$$\kappa(z) := \frac{1}{B(a,a)} z^{a-1}(1-z)^{b-1} := \frac{\pi(z)}{B(a,a)}. \tag{S1}$$

For the split scenario, we must condition on the allele frequency at the time of the split. The joint density of a single site is then,

$$
\begin{aligned}
f(z_\tau, z_0) &= \int_{z_s=0}^{1} f(z_\tau, z_0 | Z(\tau_s) = z_s)\kappa(z_s) dz_s \\
&= \int_{z_s=0}^{1} p(z_s, z_\tau; \tau - \tau_{\text{split}}) p(z_s, z_0; \tau_{\text{split}})\kappa(z_s) dz_s,
\end{aligned} \tag{S2}
$$

where $z_s$ is the integration variable for the allele frequency at the time of the split. We can further simplify Equation (S2) by twice substituting the spectral representation of the transition density (Supplementary Text S1.6),

$$
\begin{aligned}
&f(z_\tau, z_0) \\
&= \int_{z_s=0}^{1} \left( \sum_{k=0}^{\infty} \frac{e^{-\lambda_k(\tau_{\text{split}}-\tau)}}{\langle B_k, B_k\rangle_\pi} B_k(z_s) B_k(z_\tau)\pi(z_\tau) \right) \\
&\qquad \left( \sum_{m=0}^{\infty} \frac{e^{-\lambda_m \tau_{\text{split}}}}{\langle B_m, B_m\rangle_\pi} B_m(z_s) B_m(z_0)\pi(z_0) \right) \frac{\pi(z_s)}{B(a,b)} dz_s \\
&= \sum_{k=0}^{\infty} \frac{e^{-\lambda_k(2\tau_{\text{split}}-\tau)}}{\langle B_k, B_k\rangle_\pi} B_k(z_\tau) B_m(z_0) \frac{\pi(z_\tau)}{B(a,b)} \pi(z_0),
\end{aligned} \tag{S3}
$$

where we exchanged integration and summation. We recognize that this equation is equal to,

$$
f(z_\tau, z_0) = p(z_\tau, z_0; 2\tau_{\text{split}} - \tau) \frac{\pi(z_\tau)}{B(a,b)} = p(z_\tau, z_0; 2\tau_{\text{split}} - \tau)\kappa(z_s). \tag{S4}
$$

Thus, the joint density in the instance of a population split is of the same form as in the single population scenario, but with a modified time argument: $2\tau_{\text{split}} - \tau$ instead of $\tau$.

#### S1.2 Power to detect a significant association in a GWA study

We follow [1] (who follow [2]) in modeling the power of a GWA study to detect a significant association. We assume that conditional on the true effect size $\beta_\ell$, and the population allele frequency $Z_\ell$ (implicitly assuming $\hat{Z}_\ell \approx Z_\ell$) the estimated marginal effect $\hat{\beta}_\ell$ is normally distributed,

$$
\hat{\beta}_\ell | \beta_\ell, Z_\ell \sim \mathcal{N}\left(\beta_\ell, \frac{V_p}{2nZ_\ell(1-Z_\ell)}\right), \tag{S5}
$$

where $V_p$ is the total phenotypic variance which includes both genetic and environmental effects. Under the null hypothesis, i.e. $\beta_\ell = 0$, $\hat{\beta}_\ell$ is normally distributed with mean zero and the same variance as Equation (S5). The estimated contribution of locus $\ell$ to the phenotypic variance, $\hat{v}_\ell$, is $\hat{v}_\ell \coloneqq 2\hat{\beta}_\ell^2 Z_\ell(1-Z_\ell)$. When normalized by $\frac{V_p}{n}$, $\hat{v}_\ell$ is chi-squared distributed with one degree of freedom,

$$
\frac{\hat{v}_\ell}{V_p/n} = \frac{2\hat{\beta}_\ell^2 Z_\ell(1-Z_\ell)}{V_p/n} \sim \chi_1^2. \tag{S6}
$$

It follows that there is some threshold contribution to variance, $v_*$, such that the test statistic given in Equation (S6) is statistically significant. Specifically, fixing the significance threshold $\alpha$,

$$
v_* = F^{-1}(1-\alpha) = 2\left(\text{erf}^{-1}(1-\alpha)\right)^2, \tag{S7}
$$

where $F^{-1}(\cdot)$ is the inverse cumulative distribution function (*cdf*) of a chi-squared distributed random variable with one degree of freedom, and erf is the error function (eq. *A82* in [1]). This implies that a locus which satisfies,

$$\frac{2\hat{\beta}_\ell^2 Z_\ell(1-Z_\ell)}{V_p/n} > v_*, \tag{S8}$$

will yield a statistically significant association. Equation (S8) further implies that if, for a fixed $Z_\ell$,

$$|\hat{\beta}_\ell| > \sqrt{\frac{v_*(V_p/n)}{2Z_\ell(1-Z_\ell)}}, \qquad \text{or, for a fixed } \hat{\beta}, \qquad Z_\ell(1-Z_\ell) > \frac{v_*(V_p/n)}{2\hat{\beta}_\ell^2}, \tag{S9}$$

site $\ell$ will yield a significant association. If we substitute the true effect $\beta_\ell$ for $\hat{\beta}_\ell$ in Equation (S9), we can define these thresholds with respect to the true effect. And, for a fixed $\beta$ our condition is,

$$\frac{1}{2} - \frac{1}{2}\sqrt{1 - \frac{2v_*(V_p/n)}{\beta_\ell^2}} < Z_\ell < \frac{1}{2} + \frac{1}{2}\sqrt{1 - \frac{2v_*(V_p/n)}{\beta_\ell^2}}, \tag{S10}$$

We define,

$$\gamma_\ell = 1 - \sqrt{1 - \frac{2v_*(V_p/n)}{\beta_\ell^2}}, \qquad \text{and,} \qquad \beta_* = \sqrt{\frac{v_*(V_p/n)}{2Z_\ell(1-Z_\ell)}}. \tag{S11}$$

Equation (S11) specifies the threshold model used in our model, given in Equation (3), $D_\ell \in [d_\ell, 2n - d_\ell]$, where $D_\ell$ is the allele count in the GWA study and $d_\ell = \lceil n\gamma_\ell \rceil$. The distributional assumptions in Equation (S5) imply that the threshold model will be a good approximation when $n$ is large relative to $V_p$ and the detection threshold is not too small. For example, if the allele frequency in the GWA study was at its minimum frequency of $\frac{1}{2n}$, then the variance of $\hat{\beta}$ would be proportional to $V_p$—which may be large.

#### S1.3 Simulation procedures

In this section, we describe how we simulated the ancient polygenic scores (i) under neutrality and (ii) in the presence of genic selection.

**Neutrality.** To assess the accuracy of our theoretical results for the various statistics presented in Section 3, we simulated realizations of the polygenic score for ancient individuals according to our model (Section 2).

*Initialization.* To initialize each realization, we sampled $L$ population allele frequencies, $\boldsymbol{Z}(\tau) \in [0,1]^L$, from a Beta-distribution with parameters $a = b = 4N\mu$ and $b = 4N\nu$. As the population size $N$ is finite, the beta-distribution is a continuous approximation to the discrete probability mass function governing the allele frequencies. Thus, we conduct one round of binomial sampling to obtain frequencies in the set $\{0, \frac{1}{2N}, \ldots, 1 - \frac{1}{2N}, 1\}$.

*Allele frequency evolution.* Allele frequencies then evolve forward-in-time until the present ($t = 0$) when the GWA study is conducted. For forward and backward mutation rates $\mu$ and $\nu$, the transition probability of the discrete Wright-Fisher process is,

$$\psi_\mu(z) = (1-z)\mu + z(1-\nu) = \mu(1-2z) + z, \tag{S12}$$

for an allele frequency $z \in [0,1]$, and where the second equality follows for $\mu = \nu$. Conditional on the allele frequency at $t$ (generations in the past), the allele frequency in the subsequent generation is given by $(t-1)$,

$$Z_\ell(t-1)|Z_\ell(t) \sim Bin\left(2N, \psi_\mu\left(Z_\ell(t)\right)\right), \tag{S13}$$

until $t-1=0$.

*Genome-wide association study.* To conduct the GWA study, we sample $n$ diploid genotypes, $\boldsymbol{X}_i(0) \in \{-1,0,1\}^L$, for $i \in \{1,\ldots,n\}$ conditional on the allele frequencies, $\boldsymbol{Z}(0)$. Conditional on $\boldsymbol{Z}(0)$, each genotype is *iid*, $\boldsymbol{X}_i(0)|\boldsymbol{Z}(0) \sim \prod_{\ell=1}^{L} Bin(2, Z_\ell(0))$. We then sample their phenotypes, $\boldsymbol{Y}(0) \in \mathbb{R}^n$ conditional on the their genotypes, according to Equation (1). This set of $n$ genotypes and phenotype comprise the GWA study sample.

To estimate the effects we first compute the allele count at each site $D_\ell$ in the study sample. If $D_\ell$ is within the specified interval $[d_\ell, 2n-d_\ell]$, we set the effect estimate to $\beta_\ell$, as in Equation (3). If $D_\ell$ falls outside of the interval, then the effect estimate is set to 0. We then estimate $\hat{C}$ using Equation (5).

*Sampling the ancient individual(s).* We sample the genotype $\boldsymbol{X}(\tau)$ and phenotype $Y(\tau)$ of a single ancient individual conditional on the population allele frequencies $\boldsymbol{Z}(\tau)$.

*Computing estimates of the statistics.* We compute method of moments estimators for each of the statistics defined in Section 2.5. For each ancient sampling time, $\tau = \{\tau_1, \tau_2, \ldots, \tau_T\}$, we conduct $K$ simulations. For $bias(\tau)$, $mse(\tau)$, and $\hat{V}_A(\tau)$ we are interested in per-locus statistics, we average over all $L \times K$ independent locus trajectories for each time point. For example, the estimator of the *bias* is given by,

$$\overline{bias_\ell}(\tau) := \frac{1}{KL}\sum_{k=1}^{K}\sum_{\ell=1}^{L}\left(\bar{X}_{k\ell} - X_{k\ell}(\tau)\right)\left(\beta - \hat{\beta}_{k\ell}\right),$$

where $k$ indexes the simulation and $\ell$ the locus. The estimator's $(1-\alpha)\%$ confidence interval is given by,

$$bias_\ell(\tau) \in [\overline{bias_\ell}(\tau) \pm z_{\alpha/2} s_{bias_\ell}]$$

where $s_{bias_\ell}$ is the estimated standard deviations of $bias_\ell(\tau)$ and $z_{\alpha/2}$ is the inverse cumulative distribution function of a standard normally distributed random variable evaluated at $\alpha/2$.

Our estimators for the sample correlation coefficient $r^2(\tau)$ and its approximation $\rho^2(\tau)$ are computed for each replicate of $L$ loci. For example, for the $k$-th replicate,

$$r_k^2 := \frac{Cov[\hat{\boldsymbol{Y}}_k(\tau), \boldsymbol{Y}_k(\tau)]}{Var[\hat{\boldsymbol{Y}}_k(\tau)]Var[\boldsymbol{Y}_k(\tau)]} = \frac{Cov[\sum_{\ell=1}^{L}\boldsymbol{X}_{k\ell}(\tau)\hat{\beta}_{k\ell}, \sum_{\ell=1}^{L}\boldsymbol{X}_{k\ell}(\tau)\beta_{k\ell} + \boldsymbol{\epsilon}]}{Var[\sum_{\ell=1}^{L}\boldsymbol{X}_{k\ell}(\tau)\hat{\beta}_{k\ell}]Var[\sum_{\ell=1}^{L}\boldsymbol{X}_{k\ell}(\tau)\beta_{k\ell} + \boldsymbol{\epsilon}]}, \tag{S14}$$

where $\hat{\boldsymbol{Y}}_k(\tau)$ and $\boldsymbol{Y}_k(\tau)$ are the $n_a$-length vectors of ancient polygenic scores and phenotypes, respectively; $\boldsymbol{X}_{k\ell}(\tau)$ is the vector of ancient genotypes at the $\ell$-th site; and $\boldsymbol{\epsilon}$ is the vector of environmental contributions to each individual's phenotype. Our estimator $\hat{r}^2$ is an average of the $K$ realizations of $r_k^2$. To estimate $\rho^2(\tau)$, we first find estimators for the covariance and variance terms in Equation (S14) by averaging over the $K$ simulations. We then compute the ratio of these quantities to compute $\hat{\rho}^2(\tau)$.

**Genic selection.** To investigate how positive selection influences the statistical properties of polygenic scores, we simulated a recent directional selection scenario. The population evolves neutrally until the onset of selection $\tau_s$ years in the past. The $A_2$ allele confers a fitness advantage of $s$, such that the relative fitnesses of the genotypes $A_1A_1$:$A_1A_2$:$A_2A_2$ are given by $1$:$1+s$:$1+2s$.

*Initialization.* To initialize each realization, we sample $L$ population allele frequencies, $\boldsymbol{Z}(\tau) \in [0,1]^L$, from the stationary distribution of the neutral Wright-Fisher diffusion with recurrent mutation. If the ancient sampling time $\tau$ is greater than $\tau_s$ then we simulate 50 generations of neutral evolution before the onset of selection.

*Allele frequency evolution.* Allele frequencies evolve neutrally until the onset of selection at $\tau_s$. At this juncture, the allele frequencies begin to evolve according to,

$$\psi_{\mu s}(z) = \frac{\left[(1-z)^2 + z(1-z)(1+s)\right]\mu + \left[z(1-z)(1+s) + z^2(1+2s)\right](1-\mu)}{\bar{w}(z)}, \tag{S15}$$

where $\mu$ is the forward and backward per-locus, per-generation mutation rate, and the denominator is the mean fitness in a population with $A_2$ allele frequency $z$, up to the present day. The simulations with selection are otherwise identical to those under neutrality.

#### S1.4 An additional threshold model

In the main text, we introduced a simple threshold model for the effect estimates in Equation (3). Here, we consider a more realistic model in which the effect estimate $\hat{\beta}_\ell$ is the maximum-likelihood estimate (MLE) of $\beta_\ell$. We give expressions for the first two moments of $\hat{\beta}_\ell$ conditional on the contemporary allele frequencies $Z_\ell$ under each model. In doing so, we illustrate the additional challenges posed by the MLE model, and why we ultimately opted to pursue the simpler threshold model presented in the main text in Equation (3).

**Maximum-likelihood threshold model.** We define the MLE threshold model,

$$\hat{\beta}_\ell := \begin{cases} \frac{Cov[\boldsymbol{X}_\ell, \boldsymbol{Y}]}{Var[\boldsymbol{X}]} = \frac{\sum_{i=1}^n (Y_i - \bar{Y})(X_{i\ell} - \bar{X}_\ell)}{\sum_{i=1}^n (X_{i\ell} - \bar{X}_\ell)^2} & \text{if } D_\ell \in [d_\ell, 2n - d_\ell] \\ 0 & \text{else,} \end{cases} \tag{S16}$$

where each genotype, phenotype pair $(\boldsymbol{X}_i, Y_i)$ for $i \in \{1, \ldots, n\}$ is associated with an individual in the GWA study; $Cov[\cdot,\cdot]$ and $Var[\cdot]$ are the sample covariance and variance, respectively; and, as before, $\bar{X}_\ell$ and $\bar{Y}$ are the average genotype at the $\ell$-th locus and phenotype in the GWA sample.

**First moment of $\hat{\beta}$.** For both models, it can be shown that,

$$\mathbb{E}\left[\hat{\beta}_\ell | Z_\ell, \beta_\ell\right] = \beta_\ell \wp(Z_\ell, 2n, d_\ell), \tag{S17}$$

where $Z_\ell$ is the contemporary population allele frequency; and

$$\wp(z_\ell, 2n, d_\ell) = \sum_{i=d_\ell}^{2n-d_\ell} \binom{2n}{i} z_\ell^i (1-z_\ell)^{2n-i} \tag{S18}$$

is the probability that the allele count in the GWA study falls at or above the threshold $d_\ell$. Thus, when the site is segregating at a sufficiently high frequency in the GWA study sample, the estimator is unbiased. Unconditionally, for an allele count threshold $d_\ell$, $\mathbb{E}[\hat{\beta}_\ell|\beta_\ell] = \beta_\ell(1-2P^{(d_\ell)})$, where $P^{(d_\ell)}$ is the *cdf* of a beta-binomial random variable and is defined in Equation (S35).

**Second moment of $\hat{\beta}$.** The two models yield different second moments.

*Simple threshold model.* It can be shown that under the simpler model,

$$\mathbb{E}\left[\hat{\beta}_\ell^2 | \boldsymbol{Z}, \beta_\ell\right] = \beta_\ell^2 \wp(Z_\ell, 2n, d_\ell). \tag{S19}$$

As Equation (S19) only involves the allele frequency of site $\ell$, it is not influenced by variation at other sites in $\mathcal{L}$. And, unconditionally, $\mathbb{E}[\hat{\beta}_\ell^2|\beta_\ell] = \beta_\ell^2\left(1 - 2P^{(d_\ell)}\right)$.

*MLE model.* It can be shown that under the MLE model,

$$\mathbb{E}\left[\hat{\beta}_\ell^2 | \boldsymbol{Z}(0)\right] \approx \left(\beta_\ell^2 + \sum_{\ell' \neq \ell} \beta_{\ell'}^2 \frac{Z_{\ell'}(1 - Z_{\ell'})}{nZ_\ell(1 - Z_\ell)} + \frac{\sigma_e^2}{2nZ_\ell(1 - Z_\ell)}\right) \wp(Z_\ell, 2n, d_\ell), \tag{S20}$$

where the sum is over all loci $\ell' \in \mathcal{L}$ such that $\ell' \neq \ell$. The approximation is due to approximating the expectation of a ratio with the ratio of expectations, and as such, comes with all of the corresponding dangers of such an approximation. In addition, we can see from Equation (S20) that the second moment of $\hat{\beta}_\ell$ depends on the allele frequencies at all other loci in $\mathcal{L}$. While were able to compute the metrics under this approximate MLE model, we concluded that its reliance on strict assumptions about the genetic architecture (via the second moment) obscured the effects of allelic turnover. In addition, a threshold model arises naturally as the large $n$ limit of the MLE model which is, up to a sample size factor, equivalent to Equation (S5) when the allele frequency $Z_\ell$ is not too small.

#### S1.5 Polygenic scores from centered and scaled GWAS data

In the main text, we chose not to center and scale the genotypes and phenotypes of sampled individuals when conducting the GWA study. In this section, we show that our conclusions are robust to this choice. Our calculations also demonstrate that procedures convenient for statistical analysis, namely scaling, prove inconvenient when evolutionary processes are taken into account.

**Centering and scaling.** We center and scale to unit variance the phenotypes and genotypes in the GWA study,

$$\tilde{Y}_i = \frac{Y_i - \bar{Y}}{s_Y} \qquad \text{and} \qquad \tilde{X}_{i\ell} = \frac{X_{i\ell} - \bar{X}_\ell}{s_\ell}, \tag{S21}$$

where $s_Y$ and $s_\ell$ are the sample standard deviations of the phenotype and genotype at locus $\ell$, respectively. In this case, the marginal effect estimate of locus $\ell$ will be,

$$\tilde{\beta}_\ell = \frac{Cov[\tilde{\boldsymbol{X}}_\ell, \tilde{\boldsymbol{Y}}]}{Var[\tilde{\boldsymbol{X}}_\ell]} = \frac{1}{n}\sum_{i=1}^{n}(\tilde{X}_{i\ell} - 0)(\tilde{Y}_i - 0) = \frac{1}{ns_\ell s_Y}\sum_{i=1}^{n}(X_{i\ell} - \bar{X}_\ell)(Y_i - \bar{Y}) = \frac{s_\ell \hat{\beta}_\ell}{s_Y}. \tag{S22}$$

Thus, with normalized genotypes and phenotypes, the effect estimate is scaled by a factor $\frac{s_\ell}{s_Y}$, but is otherwise unaltered.

The polygenic score in the transformed case $\hat{Y}^*$, ignoring any intercept term (which would be 0), is,

$$\hat{Y}_i^* = \sum_{\ell=1}^{L} \tilde{\beta}_\ell \tilde{X}_{i\ell} = \frac{1}{s_Y}\sum_{\ell=1}^{L} \hat{\beta}_\ell (X_{i\ell} - \bar{X}_\ell). \tag{S23}$$

With centering and scaling, the polygenic score is a genetic prediction less the average genetic prediction in the GWA study sample, both scaled by a factor of $s_Y$.

**Bias.** We can compute the *bias* of the rescaled polygenic score as,

$$\begin{aligned}
\mathbb{E}\left[s_Y \hat{Y}_i^* - (Y_i - \bar{Y})\right] &= \sum_{\ell=1}^{L} \mathbb{E}\left[\hat{\beta}_\ell X_{i\ell}\right] - \sum_{\ell=1}^{L} \mathbb{E}\left[\hat{\beta}_\ell \bar{X}_\ell\right] - \mu - \sum_{\ell=1}^{L} \left[\beta_\ell X_{i\ell}\right] + \mathbb{E}\left[\bar{Y}\right] \\
&= \sum_{\ell=1}^{L} \mathbb{E}\left[\left(\hat{\beta}_\ell - \beta_\ell\right) X_{i\ell}\right] + \sum_{\ell=1}^{L} \mathbb{E}\left[\hat{\beta}_\ell \bar{X}_\ell\right] - \mu + \mu + \sum_{\ell=1}^{L} \beta_\ell \bar{X}_\ell \\
&= \sum_{\ell=1}^{L} \mathbb{E}\left[\left(\hat{\beta}_\ell - \beta_\ell\right)\left(X_{i\ell} - \bar{X}_\ell\right)\right]
\end{aligned} \tag{S24}$$

Equation (S24) shows that when we center and scale the data, we arrive at the same result. (One must rescale either $\tilde{Y}_i$ or $Y_i - \bar{Y}$ by $s_Y$ or its inverse, respectively, to put the polygenic score and the phenotype on the same scale. The former is more mathematically convenient.)

**Mean-squared error.** The proof for the *mse* is almost identical to that for the *bias*. We thereby omit it.

**Additive genetic variance.** If the effect estimates $\tilde{\boldsymbol{\beta}}$ are used instead of $\hat{\boldsymbol{\beta}}$, one must rescale the estimate of heterozygosity from the ancient sample by that estimated in the GWA study (see Equation 1 of Supplementary Note 1 of Wang et al. [3] for a related procedure),

$$\tilde{V}_A(\tau) = 2\sum_{\ell=1}^{L} \mathbb{E}\left[\tilde{\beta}_\ell^2 \left(\frac{1}{s_\ell^2}\right) \hat{Z}_\ell(\tau)\left(1 - \hat{Z}_\ell(\tau)\right)\right] = 2\sum_{\ell=1}^{L} \mathbb{E}\left[\left(\frac{1}{s_Y}\right)^2 \hat{\beta}_\ell^2 \hat{Z}_\ell(\tau)\left(1 - \hat{Z}_\ell(\tau)\right)\right]. \tag{S25}$$

This formulation is less convenient because it has random quantities in both the numerator and the denominator. Given that the units in which $Y$ is measured are arbitrary, we forewent coping with the additional complexity imposed by this scaling.

**Correlation coefficient.** By similar arguments, one can show that the sample correlation coefficient is not influenced by centering and scaling.

#### S1.6 Spectral representation of the transition density

Because the *spectral representation* of the *transition density* of an allele frequency is so central to our work, we provide a concise exposition here. For a lengthier treatment, we refer the reader to [4] and [5].

We represent the Wright-Fisher diffusion by its backward generator, $\mathscr{L}$. Introducing the quantities $a = 4N\mu$ and $b = 4N\nu$ for the population scaled mutation rates, $\mathscr{L}$ is given by,

$$\mathscr{L} f(z) = \frac{1}{2} z(1-z)\frac{\partial^2}{\partial z^2}\{f(z)\} + \frac{1}{2}[a(1-z) - bz]\frac{\partial}{\partial z}\{f(z)\}, \tag{S26}$$

where $z$ is frequency of the $A_2$ allele, and $f$ is a twice continuously differentiable bounded function on $[0, 1]$ [4].

The transition density of the Wright-Fisher diffusion, $p(z, z'; t)$, specifies the likelihood of transitioning from allele frequency $z$ to $z'$ in a time interval $[t, 0]$. The spectral representation expresses the transition density as an infinite sum,

$$p(z, z'; t) = \sum_{j=0}^{\infty} c_j(z') e^{-\lambda_j t} R_j(z), \tag{S27}$$

where, for $j = 0, 1, 2, \ldots,$ $c_j(\cdot)$ is a constant factor that depends on the initial condition, defined below in Equation (S29); $\lambda_j$ is the eigenvalue that corresponds to eigenfunction $R_j(\cdot)$; and $R_j(\cdot)$ is the $j$-th eigenfunction. The function, $\pi(\cdot)$, is the stationary measure and $\langle \cdot, \cdot \rangle_\pi$ is the inner product with respect to this measure, defined in Equation (S30). In the neutral, recurrent mutation model, the stationary measure $\pi(\cdot)$ is given by,

$$\pi(z) = z^{a-1}(1-z)^{b-1}, \tag{S28}$$

where $a$ and $b$ are the population-scaled mutation rates defined in Equation (S26). Notice that Equation (S28) is equivalent to the unnormalized density of a beta-distributed random variable with shape parameters $a$ and $b$; when normalized to integrate to one, $\pi(\cdot)$ is the stationary density $\kappa(\cdot)$, first defined in Equation (S1). When the initial condition is a point mass at the initial allele frequency, $z$, the factor $c_j(z')$ is,

$$c_j(z') = \frac{R_j(z')\pi(z')}{\langle R_j, R_j \rangle_\pi}, \tag{S29}$$

where $\langle R_j, R_j \rangle_\pi$ is the inner product of $R_j(\cdot)$ with itself. We also refer to an inner product of the form $\langle R_j, R_j \rangle$ as the squared norm of the $j$-th eigenfunction. More generally, we define the inner product of two arbitrary functions, $f(\cdot)$ and $g(\cdot)$ as,

$$\langle f, g \rangle_\pi := \int_{y=0}^{1} f(y)g(y)\pi(y)dy. \tag{S30}$$

The inner product of two eigenfunctions, $R_j$ and $R_k$, is then a special case of Equation (S30), with

$$\langle R_j, R_k \rangle_\pi = \begin{cases} \Delta_j(a,b) & \text{for } k = j, \\ 0 & \text{else,} \end{cases} \tag{S31}$$

where,

$$\Delta_j(a,b) = \frac{\Gamma(j+a)\Gamma(j+b)}{(2j+a+b-1)\Gamma(j+a+b-1)\Gamma(j+1)}, \tag{S32}$$

and $\Gamma(z) = \int_0^\infty x^{z-1}e^{-x}dx$ for $z \in \mathbb{R}$. Our work involves many inner products of the form, $\langle R_j, P_k \rangle_\pi$, where $P_k(\cdot)$ is a polynomial of degree $k$.

In the neutral recurrent mutation model, the eigenfunctions of the Wright-Fisher diffusion are Jacobi polynomials [4]. The Jacobi polynomials are polynomials of increasing order coincident with their indices, $j = 0, 1, 2, \ldots,$ and obey a three-term recurrence relation,

$$\begin{aligned} zR_j(z) = {} & \frac{(j+a-1)(j+b-1)}{(2j+a+b-1)(2j+a+b-2)} R_{j-1}(z) \\ & + \left[ \frac{1}{2} - \frac{b^2 - a^2 - 2(b-a)}{2(2j+a+b)(2j+a+b-2)} \right] R_j(z) \\ & + \frac{(j+1)(j+a+b-1)}{(2j+a+b)(2j+a+b-1)} R_{j+1}(z). \end{aligned} \tag{S33}$$

For $j = 0$,

$$zR_0(z) = \frac{a}{a+b} R_0(z) + \frac{1}{a+b} R_1(z), \tag{S34}$$

with $R_0(z) \equiv 1$.

In our work we will exploit two properties of the Jacobi polynomials: (i) the orthogonality of the eigenfunctions, i.e. Equation (S31), and (ii) the fact that a Jacobi polynomial of degree $j$ is orthogonal to all lower order polynomials (of degree $k$, $k < j$).

### S2 Extended results

We provide detailed derivations of the metrics used to characterize ancient polygenic scores. For all but $bias(\tau)$, we restrict ourselves to equal detection thresholds, although our framework readily accommodates asymmetric detection thresholds. In order to represent the metrics succinctly, we introduce several variables:

$$\begin{aligned}
P^{(d)} &:= \sum_{i=0}^{d-1} \binom{2n}{i} \frac{B(a+i, a+2n-i)}{B(a,a)} \\
P_1^{(d)} &:= \sum_{i=0}^{d-1} \left(\frac{i-n}{n}\right)^2 \binom{2n}{i} \frac{B(a+i, a+2n-i)}{B(a,a)} \\
P_2^{(d)} &:= \sum_{i=0}^{d-1} \left(\frac{(i-n)^2}{n(a+n)}\right) \binom{2n}{i} \frac{B(a+i, a+2n-i)}{B(a,a)} = \left(\frac{n}{a+n}\right) P_1^d \\
P_3^{(d)} &:= \sum_{i=0}^{d-1} \binom{2n}{i} \frac{B(a+i, a+2n-i)}{B(a,a)} \left(\frac{(2a+1)i(i-2n)+an(2n-1)}{(2a+2n+1)(a+n)}\right) \\
P_4^{(d)} &:= \sum_{i=0}^{d-1} \binom{2n}{i} \frac{B(a+i, a+2n-i)}{B(a,a)} i.
\end{aligned} \tag{S35}$$

The sums of the variables defined in Equation (S35) for $d = 2n+1$ are,

$$S = a+1, \qquad S_1 = \frac{a+n}{(1+2a)n}, \qquad S_2 = \frac{1}{2a+1}, \qquad S_3 = 0, \qquad S_4 = n. \tag{S36}$$

respectively, and with $S_3$ provided for completeness. The pervasive beta functions in Equation (S35) are a consequence of the sampling polynomial implicit in the threshold model, see Equations (3) and (S17). For example, $P^{(d)}$ is the *cdf* of a beta-binomial random variable parameterized by the number of chromosomes in the GWA study sample $2n$ and the mutation rate $a$. As we state in the main text, $\mathbb{P}\{\hat{\beta} = \beta\} = 1 - 2P^{(d)}$ when the detection thresholds are both equal to $d$. The second variable, $P_1^{(d)}$ arises from moments of the form $\mathbb{E}[\bar{X}\hat{\beta}^2]$, the expectation of the product of the mean genotype in the GWA study sample and the effect estimate. The factor $(i-n)/n$ relates the mean genotype $\bar{X}$ to the allele count $D$, i.e. $\bar{X} = (D-n)/n$. The remaining terms are less immediately interpretable; their rational is implicit in the derivations presented below and the moments provided in Supplementary Text S2.7.

#### S2.1 A form of the polygenic score bias for arbitrary thresholds

The bias of a polygenic score for an individual sampled at time $\tau$ in the past and a GWA study conducted at present is,

$$\begin{aligned}
bias(\tau) = \mathbb{E}\left[\hat{Y}(\tau) - Y(\tau)\right] &= \mathbb{E}\left[\hat{C}\right] - C + \sum_{\ell=1}^{L} \mathbb{E}\left[X_\ell(\tau)(\hat{\beta}_\ell - \beta_\ell)\right] + \mathbb{E}\left[\epsilon(\tau)\right] \\
&= \sum_{\ell=1}^{L} \beta_\ell \mathbb{E}\left[\bar{X}_\ell\right] - \mathbb{E}\left[\bar{X}_\ell \hat{\beta}_\ell\right] + \mathbb{E}\left[X_\ell(\tau)\hat{\beta}_\ell\right] - \beta_\ell \mathbb{E}\left[X_\ell(\tau)\right].
\end{aligned} \tag{S37}$$

Further simplification of Equation (S37) yields Equation (11). Using the moments derived in Supplementary Text S2.7, we can simplify Equation (11),

$$\begin{aligned} bias_\ell(\tau) &= \beta_\ell \sum_{i=d_{\ell 1}}^{2n-d_{\ell 2}} \binom{2n}{i} \frac{B(a+i, a+2n-i)}{B(a,a)} \left(e^{-a\tau}\left(\frac{i-n}{a+n}\right) - \frac{i-n}{n}\right) \\ &= \beta_\ell \left(e^{-a\tau}\left(\frac{1}{a+n}\right) - \frac{1}{n}\right)\left[n\left(P^{(d_{\ell 1})} - P^{(d_{\ell 2})}\right) - \left(P_4^{(d_{\ell 1})} - P_4^{(d_{\ell 2})}\right)\right] \\ &\approx \beta_\ell \left(e^{-a\tau} - 1\right)\left[\left(P^{(d_{\ell 1})} - P^{(d_{\ell 2})}\right) - \frac{1}{n}\left(P_4^{(d_{\ell 1})} - P_4^{(d_{\ell 2})}\right)\right], \end{aligned} \tag{S38}$$

where the last line follows for $a \ll n$. As stated in Section 3.1, for equal mutation rates and symmetric detection thresholds, $bias(\tau)$ is 0 for all $\tau$.

In Figure S2a, we plot $bias_\ell(\tau)$ in the presence of detection asymmetry for a larger range of mutation rates than presented in the main text, $a \in \{10^{-4}, 10^{-3}, 10^{-2}, 1\}$. In addition, we vary the GWA study sample size over three orders of magnitude, $n = \{10^4, 10^5, 10^6\}$ (Figure S2a). While the mutation rate $a = 1$ is not biologically plausible—this extreme illustrates features of our model that further illuminate, by contrast, the small mutation rate regime. For example, when $a = 1$ the probability of detecting a locus as significant depends heavily on the GWA study sample size $n$ (Figure S2b). Specifically, for $a = 1$, $P^{(d_\ell)} = \frac{d_\ell}{2n}$ increases linearly with $d_\ell$ (Figure S2b; bottom). In contrast, for $a \ll 1$ and modest $n$, this probability is insensitive to $n$ (Figure S2b). Specifically, $P^{(d_\ell)} \approx 0.5$ for all values of $n$ and $d_\ell$ (Figure S2b; top) as most of the allele frequencies are very close to, or equal to zero or one, and thus will always elude detection in GWA studies with finite sample sizes. In other words, once $n$ is large enough, varying $d_\ell$ yields diminishing returns.

In Figure S2a, we set $d_{\ell 1} = 1$ and $d_{\ell 2} = n$ for each sample size to illustrate the effects of an extreme imbalance. For $d_{\ell 2} = n$, positive effect alleles cannot be detected, while $d_{\ell 1} = 1$ implies that a negative effect allele will be detected as long as it is segregating the in the GWA study sample. As $d_{\ell 1} < d_{\ell 2}$, sites where the trait-decreasing allele is at higher frequency in the GWA study ($D_\ell < n$) will be detected more often than sites where the trait-increasing allele is at higher frequency ($D_\ell > n$). This implies that sites where the majority of individuals in the GWA study possess negative effect alleles are more likely to have non-zero effects in the genetic prediction. At the same time, the majority of sites contributing to the estimated intercept $\hat{C}$ will have $\bar{X}_\ell > 0$, and thus, in expectation, $\hat{C} \geq 0$. Thus, at $\tau = 0$, the excess positive contributions to the estimated intercept are tempered by the excess negative contributions to the genetic prediction. As $\tau$ increases, $bias_\ell(\tau)$ becomes more positive (Figure S2a). This is because the estimated intercept $\hat{C}$ is constant, whereas, the expected value of the genetic prediction approaches zero with increasing $\tau$. The latter follows from the fact that as $\tau$ increases genotype of the ancient sample $X_\ell(\tau)$ becomes independent of average genotype in the GWA study $\bar{X}_\ell$, and its expected value approaches zero. Thus, in the large $\tau$ limit, $bias(\tau) = \mathbb{E}[\hat{C}]$, which is positive for $d_1 < d_2$.

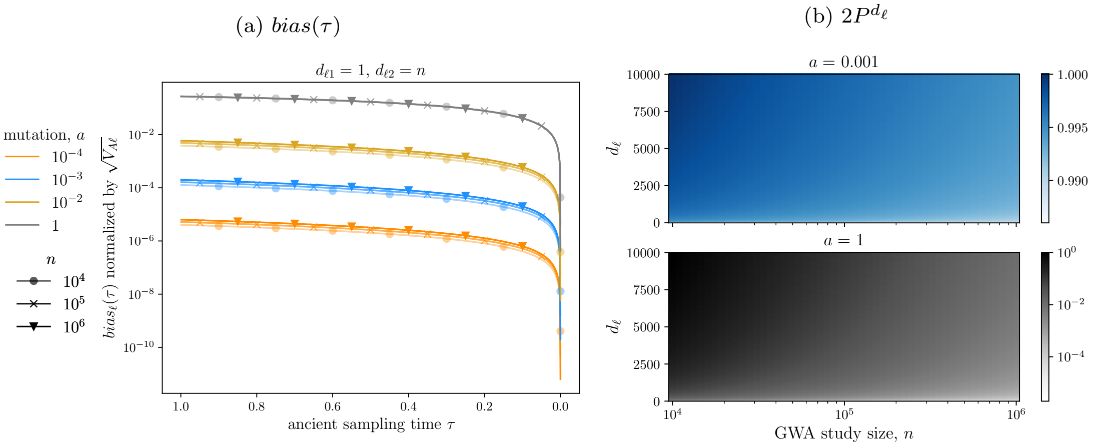

Figure S2: **Asymmetry in the detection threshold.** In (a), we plot $bias_\ell(\tau)$ for $a = 10^{-3}$ (orange) and $a = 1$ (blue) across three GWA study sizes, $n = 10^3, 10^4, 10^5$. We set the detection thresholds to $d_1 = 1$ and $d_2 = 100$. In (b), we plot $2P^{d_\ell}$ as a function of $n$ and the detection threshold $d_\ell$ for $a = 10^{-3}$ (top; orange) and $a = 1$ (bottom; blue).

Approximating the increase of $bias_\ell(\tau)$, given in Equation (S38), for small $a$ and large $n$,

$$bias_\ell(\tau) \approx bias_\ell(0) + \beta_\ell a \tau \left( P^{(d_{\ell 1})} - P^{(d_{\ell 2})} \right), \tag{S39}$$

gives us additional insight into these results. In Equation (S39), $bias_\ell(0)$ is an exact expression for the $bias_\ell(\tau)$ evaluated at $\tau = 0$; and $P^{(d_{\ell i})}$ is the probability that the allele count in the GWA study, $D_\ell$, is less than $d_{\ell i}$ for $i = 1, 2$. Under our assumptions, $P^{(d_{\ell i})}$ is the cumulative distribution function (*cdf*) of a beta-binomial random variable with $2n$ trials, parameterized by the mutation rate $a$. At $\tau = 0$, the focal individual is an independent sample from the GWA study population. Thus, the intercept term captures contributes to $bias_\ell(\tau)$ exclusively due to finite sampling. For $\tau > 0$, allelic turnover induces changes in the frequencies of sites not detected in the GWA study ($\ell$ such that $\hat{\beta}_\ell = 0$), which may contribute to the phenotypic variation of ancient individuals. Thus, the linear term captures additional bias due to finite sampling *and* allelic turnover.

We conclude that increases in $bias_\ell(\tau)$ with $\tau$ depend primarily on the difference in the detection probabilities for trait-increasing and decreasing alleles, i.e. $P^{(d_{\ell 1})} - P^{(d_{\ell 2})}$. For small $a$, this difference is small relative to the (square root) of the additive genetic variance $V_A$ due to the fact that the detection probability is insensitive to the threshold $d_\ell$. However, differences in sample size are apparent when the mutation rate is small—with larger sample sizes yielding a larger bias (Figure S2a). This sample size dependency is due to the fact that increased power to detect low frequency alleles with larger $n$ results in a larger difference between the one-sided detection threshold. As $a$ approaches one, the effects of sample size diminish (in log scale). For $a = 1$, the difference in one-sided detection probabilities $P^{(1)} - P^{(n)} = \frac{n-1}{2n}$, which will be close to $\frac{1}{2}$ for modest values of $n$. In addition, for large $a$, $bias_\ell(\tau)$ is non-negligible relative to (the square root of) $\mathbb{E}[V_A]$.

#### S2.2 Deriving the mean-squared error

Substituting the definitions of $\hat{Y}(\tau)$ and $Y(\tau)$, we can simplify the expression for the mean-squared error ($mse$),

$$
\begin{aligned}
mse(\tau) = \mathbb{E}\left[\left(\hat{Y}(\tau) - Y(\tau)\right)^2\right] &= \mathbb{E}\left[\left((\hat{C} - C) + \sum_{\ell=1}^{L} X_\ell(\hat{\beta}_\ell - \beta_\ell) - \epsilon(\tau)\right)^2\right] \\
&= \mathbb{E}\left[\left(\sum_{\ell=1}^{L}(X_\ell(\tau) - \bar{X}_\ell)(\hat{\beta}_\ell - \beta_\ell) + (\bar{\epsilon} - \epsilon(\tau))\right)^2\right] \qquad \text{(S40)} \\
&= \sum_{\ell=1}^{L} \mathbb{E}\left[(X_\ell(\tau) - \bar{X}_\ell)^2(\hat{\beta}_\ell - \beta_\ell)^2\right] + \mathbb{E}\left[(\bar{\epsilon} - \epsilon(\tau))^2\right],
\end{aligned}
$$

where the cross-terms in Equation (S40) cancel due to independence between the environmental noise, which has mean 0, and the genotypes. The error term simplifies,

$$
\mathbb{E}\left[(\bar{\epsilon} - \epsilon(\tau))^2\right] = \mathbb{E}\left[\bar{\epsilon}^2 - 2\bar{\epsilon}\epsilon(\tau) + (\epsilon(\tau))^2\right] = \left(\frac{n-1}{n}\right)\sigma_e^2. \qquad \text{(S41)}
$$

When $\sigma_e^2 = 0$, the $mse$ reduces to,

$$
\begin{aligned}
mse_\ell(\tau) = \; & \mathbb{E}\left[X_\ell^2(\tau)\hat{\beta}_\ell^2\right] - 2\beta_\ell \mathbb{E}\left[X_\ell^2(\tau)\hat{\beta}_\ell\right] + \beta_\ell^2 \mathbb{E}\left[X_\ell^2(\tau)\right] \\
& - 2\left(\mathbb{E}\left[X_\ell(\tau)\bar{X}_\ell\hat{\beta}_\ell^2\right] - 2\beta_\ell \mathbb{E}\left[X_\ell(\tau)\bar{X}_\ell\hat{\beta}_\ell\right] + \beta_\ell^2 \mathbb{E}\left[X_\ell(\tau)\bar{X}_\ell\right]\right) \qquad \text{(S42)} \\
& + \mathbb{E}\left[\bar{X}_\ell^2\hat{\beta}_\ell^2\right] - 2\beta_\ell \mathbb{E}\left[\bar{X}_\ell^2\hat{\beta}_\ell\right] + \beta_\ell^2 \mathbb{E}\left[\bar{X}_\ell^2\right].
\end{aligned}
$$

For equal detection thresholds $d_{\ell 1} = d_{\ell 2} = d_\ell$, Equation (S42) reduces to,

$$
mse(\tau) = 2\sum_{\ell=1}^{L} \beta_\ell^2 \left[\left(\frac{a+1}{2a+1}\right) P^{(d_\ell)} + P_1^{(d_\ell)} - 2e^{-a\tau} P_2^{(d_\ell)} + \left(\frac{1}{2a+1}\right) e^{-(2a+1)\tau} P_3^{(d_\ell)}\right], \qquad \text{(S43)}
$$

where $d_\ell = \lceil n\gamma_\ell \rceil$. The change in $mse(\tau)$ is due to the difference between the two exponential terms in Equation (S43). From Equation (S43), we derive the derivative of $mse(\tau)$,

$$
\frac{dmse(\tau)}{d\tau} = 2\sum_{\ell=1}^{L} \beta_\ell^2 \left[2aP_2^{(d_\ell)} e^{-a\tau} - P_3^{(d_\ell)} e^{-(2a+1)\tau}\right], \qquad \text{(S44)}
$$

which, for small $a$ and $\tau$, is,

$$
\frac{dmse(\tau)}{d\tau} \approx 2\sum_{\ell=1}^{L} \beta_\ell^2 \left[2aP_2^{(d_\ell)} - P_3^{(d_\ell)} e^{-\tau}\right] \approx 2a \sum_{\ell=1}^{L} \beta_\ell^2 P^{(d_\ell)} \left(2 - e^{-\tau}\right). \qquad \text{(S45)}
$$

#### S2.3 Deriving the expected additive genetic variance

We solve for $\hat{V}_A(\tau)$ in an ancient sample of size $n_a$ and a GWA study sample of size $n$. Considering a single locus $\ell$ and conditioning on the ancient and contemporary allele frequencies,

$$\begin{aligned}\hat{V}_{A\ell}(\tau) &= 2\mathbb{E}\left[\mathbb{E}\left[\hat{\beta}_\ell^2 \hat{Z}_\ell(\tau)(1-\hat{Z}_\ell(\tau))|Z_\ell, Z_\ell(\tau)\right]\right] \\ &= 2\mathbb{E}\left[\mathbb{E}\left[\hat{Z}_\ell(\tau)(1-\hat{Z}_\ell(\tau))|Z_\ell(\tau)\right]\mathbb{E}\left[\hat{\beta}_\ell^2|Z_\ell\right]\right] \\ &= 2\left(\frac{2n_a-1}{2n_a}\right)\beta_\ell^2 \mathbb{E}\left[Z_\ell(\tau)(1-Z_\ell(\tau))\wp(Z_\ell, d_\ell, n)\right],\end{aligned} \tag{S46}$$

where $\wp(Z_\ell, 2n, d_\ell)$ is the Binomial sampling probability defined in Equation (S18). Substituting the spectral representation of the *tdf* yields,

$$\hat{V}_{A\ell}(\tau) = \beta_\ell^2\left(\frac{2n_a-1}{2n_a}\right)\left(\frac{1}{2a+1}\right)\left[a(1-2P^{(d_\ell)}) + 2e^{-(2a+1)\tau}P_3^{(d_\ell)}\right]. \tag{S47}$$

To compare $\hat{V}_A$ across parameter regimes, we normalize by the expected population additive genetic variance $\mathbb{E}[V_A]$. At stationarity and for mutation rate $a$,

$$\mathbb{E}\left[V_{A\ell}(\tau)\right] = 2\beta_\ell^2\int_{z_\tau} z_\tau(1-z_\tau)\kappa(z_\tau)dz_\tau = \left(\frac{a}{2a+1}\right)\beta_\ell^2, \tag{S48}$$

where $\kappa(\cdot)$ is the stationary density given in Equation (S1).

For small $a$, $\hat{V}_A(\tau)$ will change at rate,

$$\frac{d\mathbb{E}[\hat{V}_A(\tau)]}{d\tau} \approx -2\left(\frac{2n_a-1}{2n_a}\right)e^{-\tau}\sum_{\ell=1}^{L}\beta_\ell^2 P_3^{(d_\ell)} \approx -2\left(\frac{2n_a-1}{2n_a}\right)e^{-\tau}a\sum_{i=1}^{L}\beta_\ell^2 P^{(d_\ell)}, \tag{S49}$$

where the right hand expression follows from the approximation $P_3^{(d_\ell)} \approx aP^{(d_\ell)}$ (see Supplementary Text S2.5).

#### S2.4 Deriving the approximate sample correlation coefficient

In this subsection, we (i) describe the two approximation steps implicit in our definition of $\rho^2(\tau)$ given in Equation (10); (ii) derive an explicit form for $\rho^2(\tau)$; and (iii) derive the approximate decay of relative accuracy $\rho^2(\tau)/\rho^2(0)$.

(i) A practitioner is often interested in the accuracy of their predictor with respect to a particular sample. This sample correlation coefficient (for $n_a$ ancient individuals from $\tau$) is defined as,

$$r(\tau) := \frac{Cov[\hat{\boldsymbol{Y}}(\tau), \boldsymbol{Y}(\tau)]}{\sqrt{Var[\hat{\boldsymbol{Y}}]Var[\boldsymbol{Y}]}} = \frac{\sum_{i=1}^{n_a}(\hat{Y}_i(\tau)-\bar{\hat{Y}}(\tau))(Y_i(\tau)-\bar{Y}(\tau))}{\sqrt{\sum_{i=1}^{n_a}(\hat{Y}_i(\tau)-\bar{\hat{Y}}(\tau))^2}\sqrt{\sum_{i=1}^{n_a}(Y_i(\tau)-\bar{Y}(\tau))^2}}, \tag{S50}$$

where $Cov[\cdot,\cdot]$ and $Var[\cdot]$ are the sample covariance and variance operators, respectively; and $\hat{\boldsymbol{Y}}(\tau), \boldsymbol{Y}(\tau) \in \mathbb{R}^{n_a}$ are the $n_a$-dimensional vectors of polygenic scores and phenotypes, respectively. Ultimately, we will approximate the expectation of the squared sample correlation coefficient $r^2(\tau)$ with a ratio of expectations,

$$\mathbb{E}\left[r^2(\tau)\right] \approx \frac{\mathbb{E}\left[Cov[\hat{\boldsymbol{Y}}(\tau), \boldsymbol{Y}(\tau)]\right]^2}{\mathbb{E}[Var[\hat{\boldsymbol{Y}}(\tau)]]\mathbb{E}\left[Var[\boldsymbol{Y}(\tau)]\right]}, \tag{S51}$$

which we defined as $\rho^2(\tau)$ in Equation (10). To arrive at this approximation, we must first approximate the expectation of the ratio in Equation (S50) as the ratio of expectations. Second, we must pull the expectation inside the square roots in the denominator. A full investigation of the validity of these steps in general is beyond the scope of the present study. Rather, we validate these approximation steps by simulations of a few parameter regimes of particular interest.

After these two approximation steps, we compute the quantity in Equation (S51) exactly under our framework. We take each element of Equation (S51) in turn.

(ii) When we plug in our modeling assumptions, the numerator of Equation (S50) becomes

$$Cov[\hat{\boldsymbol{Y}}(\tau), \boldsymbol{Y}(\tau)] = \sum_{i=1}^{n_a} \sum_{\ell,\ell'} \hat{\beta}_\ell \beta_{\ell'} (X_{i\ell}(\tau) - \bar{X}_\ell(\tau))(X_{i\ell'}(\tau) - \bar{X}_{\ell'}(\tau)) + Cov[\hat{\boldsymbol{Y}}(\tau), \boldsymbol{\epsilon}], \tag{S52}$$

due to the linearity of the covariance operator, with $\boldsymbol{\epsilon} \in \mathbb{R}_a^n$ as the vector of environmental effects. In expectation, assuming *iid* loci with equal effects $\beta$ and *iid* ancient samples,

$$\mathbb{E}\left[Cov[\hat{\boldsymbol{Y}}(\tau), \boldsymbol{Y}(\tau)]\right] = \frac{1}{n_a} \sum_{i=1}^{n_a} \sum_{\ell=1}^{L} \beta_\ell \mathbb{E}\left[\hat{\beta}_\ell (X_{i\ell}(\tau) - \bar{X}_\ell(\tau))^2\right] = L\beta \mathbb{E}\left[\hat{\beta}(X_i(\tau) - \bar{X}(\tau))^2\right]. \tag{S53}$$

Similarly,

$$\mathbb{E}\left[Var[\hat{\boldsymbol{Y}}(\tau)]\right] = L\mathbb{E}\left[\hat{\beta}^2 (X_i(\tau) - \bar{X}(\tau))^2\right], \tag{S54}$$

which, under our simple threshold model is equal to the expectation of the covariance given in Equation (S53). Finally,

$$\mathbb{E}\left[Var[\boldsymbol{Y}(\tau)]\right] = L\beta^2 \mathbb{E}\left[(X(\tau) - \bar{X}(\tau))^2\right] + \left(\frac{n_a - 1}{n_a}\right) \sigma_e^2. \tag{S55}$$

All together, our approximation for $r^2(\tau)$ reduces to,

$$\mathbb{E}\left[r^2\right] \approx \frac{L\beta \mathbb{E}\left[\hat{\beta}(X(\tau) - \bar{X}(\tau))^2\right]}{L\beta^2 \mathbb{E}\left[(X(\tau) - \bar{X}(\tau))^2\right] + \left(\frac{n_a-1}{n_a}\right)\sigma_e^2} = \frac{\mathbb{E}\left[\hat{\beta}(X(\tau) - \bar{X}(\tau))^2\right]/\beta}{\mathbb{E}\left[(X(\tau) - \bar{X}(\tau))^2\right] + \left(\frac{n_a-1}{n_a}\right)\sigma_{e'}^2}, \tag{S56}$$

where $\sigma_{e'}^2 = \sigma_e^2/(L\beta^2)$. All that remains is to solve for the expectations in the numerator and denominator of Equation (S56). The first involves both the GWA study and ancient sample times,

$$\begin{aligned} \mathbb{E}\left[\hat{\beta}(X(\tau) - \bar{X}(\tau))^2\right] &= \mathbb{E}\left[\mathbb{E}\left[\hat{\beta}(X(\tau) - \bar{X}(\tau))^2 | Z(0), Z(\tau)\right]\right] \\ &= \beta \mathbb{E}\left[\wp(Z(0)) \mathbb{E}\left[(X(\tau) - \bar{X}(\tau))^2 | Z(\tau)\right]\right] \\ &= 2\beta \left(\frac{n_a - 1}{n_a}\right) \mathbb{E}\left[\wp(Z(0)) Z(\tau)(1 - Z(\tau))\right], \end{aligned} \tag{S57}$$

which we recognize as closely related to the expected estimated additive genetic variance, $\hat{V}_A(\tau)$, with $\wp(Z(0))$ defined in Equation (S18). The expectation in the denominator is,

$$\mathbb{E}\left[\hat{\beta}(X(\tau) - \bar{X}(\tau))^2\right] = 2\left(\frac{n_a - 1}{n_a}\right) \mathbb{E}\left[Z(\tau)(1 - Z(\tau))\right] = \left(\frac{n_a - 1}{n_a}\right)\left(\frac{a}{2a+1}\right), \tag{S58}$$

which is equal to the $\mathbb{E}[V_A]$ at stationarity normalized by the squared true effect $\beta^2$ and multiplied by the $n_a$-dependent factor to account for ancient sample size. We then see that our approximation to the expectation of $r^2(\tau)$ is insensitive to the ancient sample size and equal to,

$$\mathbb{E}\left[r^2(\tau)\right] \approx \rho^2(\tau) := \frac{2\mathbb{E}\left[\wp(Z(0))Z(\tau)(1-Z(\tau))\right]}{\frac{a}{2a+1}+\sigma^2_{e'}} = \left(\frac{2n_a}{2n_a-1}\right)\frac{\hat{V}_{A\ell}(\tau)/\beta^2}{\frac{a}{2a+1}+\sigma^2_{e'}}, \tag{S59}$$

where the ancient sample size dependent factor in the rightmost expression cancels with its inverse in $\hat{V}_{A\ell}(\tau)$. Thus, the sample correlation coefficient is proportional to the estimated additive genetic variance, and its derivative is given by,

$$\begin{aligned}\frac{d\mathbb{E}\left[r^2(\tau)\right]}{d\tau} &\approx \frac{d}{d\tau}\left(\frac{2n_a}{2n_a-1}\right)\frac{\hat{V}_{A\ell}(\tau)/\beta^2}{\frac{a}{2a+1}+\sigma^2_{e'}} \\ &\approx \left(\frac{1}{\frac{a}{2a+1}+\sigma^2_{e'}}\right)\left(\frac{2}{2a+1}\right)(-(2a+1))e^{-(2a+1)\tau}P_3^{(d)} \\ &\approx \left(\frac{1}{a+\sigma^2_{e'}}\right)2e^{-\tau}P_3^{(d)} \approx \left(\frac{a}{a+\sigma^2_{e'}}\right)2e^{-\tau}P^{(d)},\end{aligned} \tag{S60}$$

where the last line follows for $a \ll 1$, as $e^{-(2a+1)\tau} \approx e^{-\tau}$ and $P_3^{(d)} \approx aP^{(d)}$, see Equation (S69).

(iii) We show that for small mutation rates, relative accuracy decays at a rate that is independent of the mutation rate $a$ and detection threshold $d$. For *iid* loci,

$$\rho^2(\tau)/\rho^2(0) = \frac{a(1-2P^{(d)}) + 2e^{-(2a+1)\tau}P_3^{(d)}}{a(1-2P^{(d)}) + 2P_3^{(d)}} \approx \frac{2e^{-\tau}P_3^{(d)}}{2P_3^{(d)}} = e^{-\tau}, \tag{S61}$$

where, we have claimed that $a(1-2P^{(d)}) \approx 0$ for all $d \in \{1,\dots,n\}$, and that $2a+1 \approx 1$. If we relax the *iid* assumption, we have,

$$\rho^2(\tau)/\rho^2(0) = \frac{\sum_{\ell=1}^{L}\beta_\ell^2\left[a(1-2P^{(d_\ell)}) + 2e^{-(2a+1)\tau}P_3^{(d_\ell)}\right]}{\sum_{\ell=1}^{L}\beta_\ell^2\left[a(1-2P^{(d_\ell)}) + 2P_3^{(d_\ell)}\right]}, \tag{S62}$$

which could be computed for a given distribution of $\beta_\ell$. For $a \ll 1$, the $a(1-2P^{(d_\ell)})$ terms in Equation (S62) *may be* negligible, yielding the same result as Equation (S61), which implies that relative accuracy is insensitive to distributional assumptions on $\beta$ for small $a$. However, more rigorous theoretical and simulation-based work is required to assess the accuracy of this claim.

#### S2.5 Deriving approximations to the metrics

In the main text, we present several approximations for the initial rate of increase or decrease of the metrics. Here, we show how we arrived at these approximations from the exact forms given in the previous sections. For a given metric, we first compute a first order Taylor series expansion (in $\tau$). We then find the intercept, i.e. the value of the statistic at zero, and the slope. We subsequently make use of the following approximations: (i) $P_1^{(d)} \approx P^{(d)}$; (ii) $P_2^{(d)} \approx P^{(d)}$; and (iii) $P_3^{(d)} \approx aP^{(d)}$.

**Approximate metrics.** For the *bias*, this approach yields,

$$\begin{aligned}bias_\ell(\tau) &\approx \beta_\ell\left[(1-a\tau)\left(\frac{1}{a+n}\right) - \frac{1}{n}\right]\left[n\left(P^{(d_{\ell 1})} - P^{(d_{\ell 2})}\right) + \left(P_4^{(d_{\ell 2})} - P_4^{(d_{\ell 1})}\right)\right] \\ &\approx bias_\ell(0) + \beta_\ell \cdot a\tau\left(P^{(d_{\ell 1})} - P^{(d_{\ell 2})}\right),\end{aligned} \tag{S63}$$

where the last line follows from (i) using the approximations noted in the prelude; (ii) ignoring the $P_4^{(d_{\ell i})}$ terms in the slope (which are order $\mathcal{O}(\frac{1}{n})$); (iii) and $\frac{1}{a+n} \approx \frac{1}{n}$. Using the same approach, $mse_\ell(\tau)$ with equal thresholds $d_\ell$, becomes,

$$\begin{aligned} mse_\ell(\tau) &\approx 2\beta_\ell^2 \Big[ \left(\frac{a+1}{2a+1}\right) P^{(d_\ell)} + P_1^{(d_\ell)} - 2P_2^{(d_\ell)} + \left(\frac{1}{2a+1}\right) P_3^{(d_\ell)} + 2a\tau P_2^{(d_\ell)} - \tau P_3^{(d_\ell)} \Big] \\ &\approx mse_\ell(0) + 2\beta_\ell^2 a\tau P^{(d_\ell)}. \end{aligned} \tag{S64}$$

And, we can approximate $\hat{V}_A(\tau)$ as,

$$\begin{aligned} \hat{V}_{A\ell}(\tau) &\approx \left(\frac{2n_a-1}{2n_a}\right) \beta_\ell^2 \left(\frac{1}{2a+1}\right) \left[ \left( a(1-2P^{(d_\ell)}\right) + \left( 2\left(1-(2a+1)\tau\right) P_3^{(d_\ell)} \right) \right] \\ &\approx \hat{V}_{A\ell}(0) - 2\left(\frac{2n_a-1}{2n_a}\right) \beta_\ell^2 a P^{(d_\ell)} \tau. \end{aligned} \tag{S65}$$

The approximation for the accuracy $\rho^2(\tau)$ follows immediately from Equation (S65).

In Figure S3a,d, we plot the approximate rate of change $2aP^{(d)}$, for low and high detection thresholds. In addition, in Figure S4, we compare our exact theoretical results to their approx-

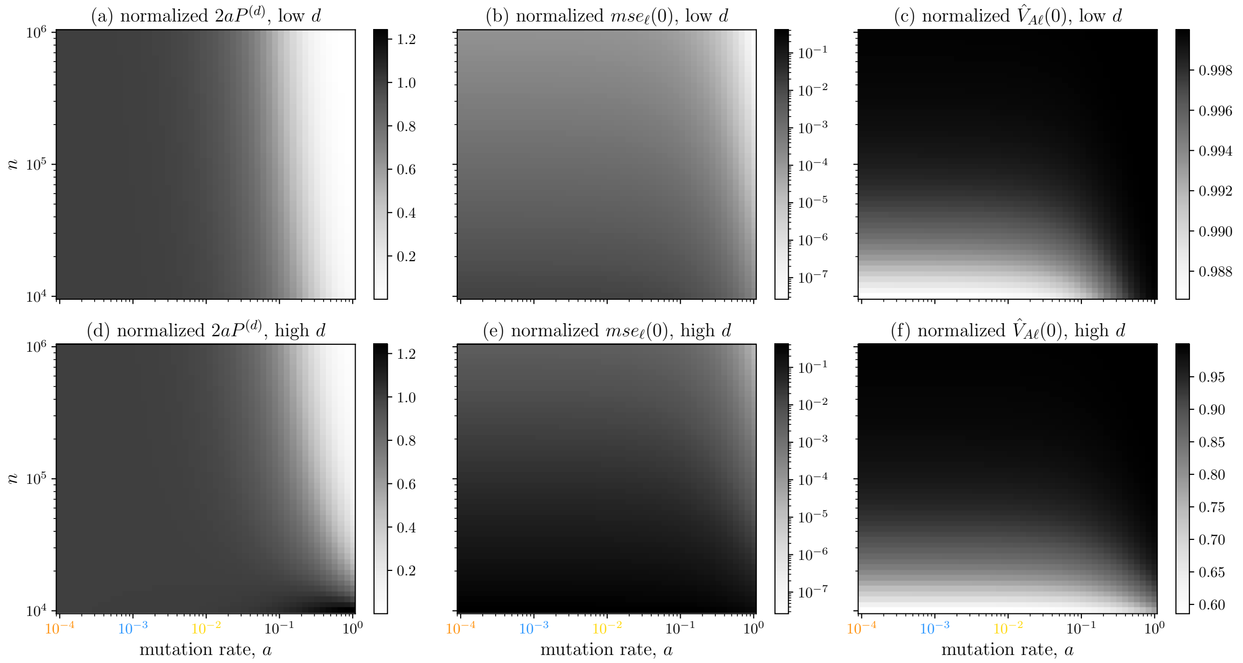

Figure S3: **Pieces of the approximation to the mean-squared error and expected estimated additive genetic variance.** In (a), we plot $2aP^{(d)}$ normalized by $\mathbb{E}[V_A] = \beta^2(\frac{a}{2a+1})$ across a range of mutation rates $a \in \{10^{-4}, \ldots, 1\}$ and GWA study sample sizes $n$, for a small detection threshold. Here, $d$ is either 132 or 133, corresponding to a squared effect size of $\beta^2 = 0.25$, when the significance threshold is $\alpha = 10^{-8}$ and the phenotypic variance $V_p = 1$. In (b) and (c), we plot the initial $mse_\ell(\tau)$ and $\hat{V}_{A\ell}(\tau)$ (both normalized by the true $V_A$) for small $d$. In (d-f), we repeat plots (a-c), except with a higher detection threshold, respectively. The smaller effect size of $\beta^2 = 0.01$ yields thresholds in the range $d \in \{3209, \ldots, 4142\}$, in order of increasing sample size. Note that in contrast to the other pairs of plots, (c) and (f) do not share a scale.

imations over a short time scale of $\tau \in [0.2, 0]$. We observe that the approximation fares better for smaller values of $d$, as well as, for larger $n$. Below, we show how some of the steps in our approximations are adversely affected by large $d$ and small $n$.

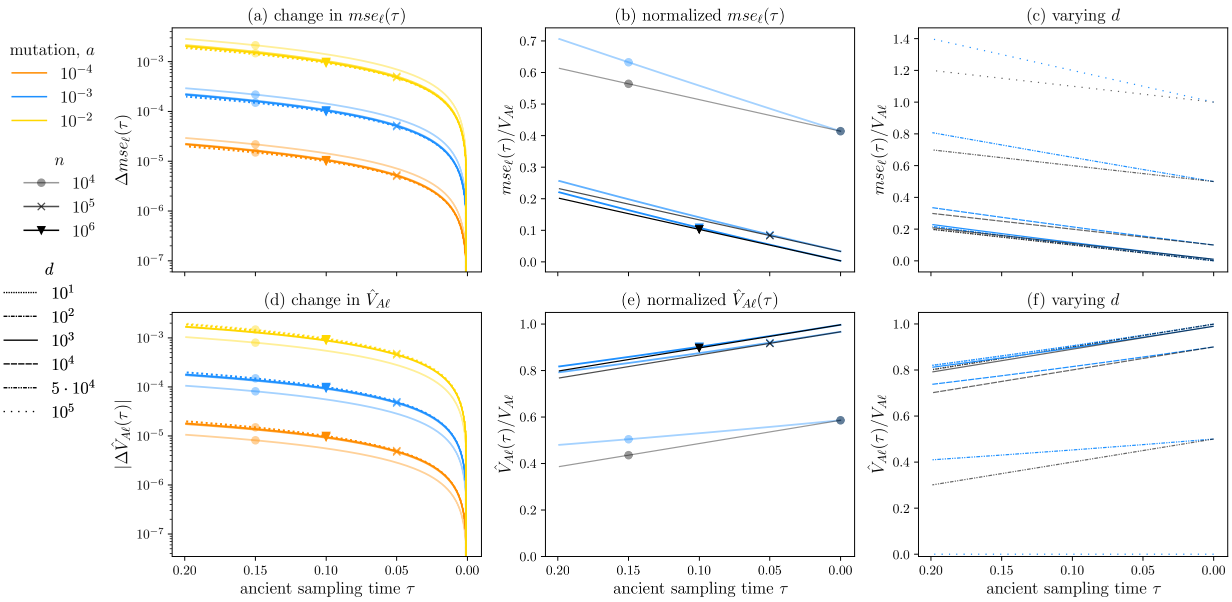

Figure S4: **Approximations for the per locus contributions to the mean-squared error and estimated additive genetic variance across sample sizes, mutation rates, and detection thresholds.** This plot is identical to Figure 2 except that we (i) include our approximations to the two statistics, Equation (15) and Equation (17), and (ii) plot our results over a short time frame, $\tau \in [0.2, 0]$. In (a), the approximations are depicted as colored, dotted lines corresponding to each mutation rate and sample size pair. In (b), the approximations are denoted by the same markers and opacity as their blue counterparts. And, in (c), the approximations are provided in black, with line pattern indicating the threshold $d$.

**Approximation error.** We quantify the error incurred in the approximation $P_1^{(d)} \approx P_3^{(d)}$ and $P_2^{(d)} \approx P_3^{(d)}$. To do so, we first express $P_1^{(d)}$ and $P_2^{(d)}$ as functions of $P_3^{(d)}$. We refer to the $i$-th term in quantities specified in Equation (S35) as $P_\cdot^i$, dropping the superscript $d$ for succinctness.

$$P_1^i = \left(\frac{i-n}{n}\right)^2 \binom{2n}{i} \frac{B(a+i, a+2n-i)}{B(a,a)} = \left(\frac{i-n}{n}\right)^2 P^i = \left(\frac{i}{n}\right)^2 P^i - 2\left(\frac{in}{n^2}\right) + P^i. \tag{S66}$$

Thus,

$$P_1^{(d)} = P^{(d)} + \frac{1}{n^2}\sum_{i=0}^{d-1} i^2 P^i - \frac{2}{n}\sum_{i=0}^{d-1} iP^i. \tag{S67}$$

Note that the summations in Equation (S67) are the second and first moments of a beta-binomial random variable truncated at $d-1$, respectively. As long as the two summations are $O(n)$ and $O(1)$, respectively, the error of the approximation will be smaller than $O(\frac{1}{n})$ as both summations are non-negative. We can repeat the same procedure for $P_2^{(d)}$,

$$P_2^{(d)} = \left(\frac{n}{a+n}\right) P^{(d)} + \frac{1}{n(a+n)}\sum_{i=0}^{d-1} i^2 P^i - \left(\frac{2}{a+n}\right)\sum_{i=0}^{d-1} iP^i \approx P_1^{(d)}, \tag{S68}$$

where the approximation is valid for $a \ll n$, and in this regime, our analysis in the previous paragraph also applies to $P_2^{(d)}$.

Finally, we consider the approximation $P_3^{(d)} \approx aP^{(d)}$,

$$\begin{aligned}
P_3^i &= P^i\left(\frac{(2a+1)i(i-2n)+an(2n-1)}{(2a+2n+1)(a+n)}\right)+aP^i-aP^i\\
&= aP^i+P^i\left(\frac{(2a+1)i(i-2n)+an(2n-1)-a(2a+2n+1)(a+n)}{(2a+2n+1)(a+n)}\right)\\
&= aP^i+P^i\left(\frac{(2a+1)i(i-2n)-a(2a^2+4an+a+2n)}{(2a+2n+1)(a+n)}\right)\\
&= aP^i+P^i\left(\frac{(2a+1)i(i-2n)-a(2a+1)(a+2n)}{(2a+2n+1)(a+n)}\right)\\
&= aP^i+(2a+1)P^i\left(\frac{i(i-2n)-a(a+2n)}{(2a+2n+1)(a+n)}\right).
\end{aligned} \tag{S69}$$

Thus,

$$P_3^{(d)} = aP^{(d)} - \frac{(2a+1)}{(2a+2n+1)(a+n)}\left[a(a+2n)(d-1)P^{(d)}+\sum_{i=0}^{d-1}\binom{2n}{i}\frac{B(a+i,a+2n-i)}{B(a,a)}i(2n-i)\right], \tag{S70}$$

And,

$$|P_3^{(d)} - aP^{(d)}| \approx \frac{a(d-1)}{n}P^{(d)}+\sum_{i=0}^{d-1}\binom{2n}{i}\frac{B(a+i,a+2n-i)}{B(a,a)}\left(\frac{i}{n}-\frac{i^2}{2n^2}\right) \tag{S71}$$

where the approximation follows for $a \ll 1$ and $a \ll n$. Thus, $P_3^{(d_\ell)} \approx aP^{(d_\ell)}$ will be a very good approximation when $d \ll n$, but should also hold for modest $d$ as long $n$ is reasonably large. It is possible that when the mutational target is very large, e.g. $O(n)$, the approximation errors may be non-negligible for large enough $d$. However, large $d$ implies a small $\beta$, thereby tempering any approximation errors in practice. An additional benefit of expressing $P_1^{(d)}$, $P_2^{(d)}$, and $P_3^{(d)}$ in terms of $P^{(d)}$ is that we can now take advantage of efficient coding of the beta-binomial probability mass function in the Python module `scipy` to compute analytical results for larger values of $d$.

**Computations for large $n$ and $d$.** For large $n$, computing the terms in Equation (S35), excluding $P^{(d)}$ (which we compute using `scipy`), becomes computationally prohibitive. However, for large $n$, we can approximate these quantities as follows. Defining $z = \frac{d}{2n}$ and the incomplete beta function as $I_z(x,y) = \frac{1}{B(x,y)}\int_0^z z^{x-1}(1-z)^{y-1}dz$,

$$\begin{aligned}
P_1^{(d)} &\approx \left(\frac{1}{n^2}\right)I_z(a+2,a)-\left(\frac{2}{n}\right)I_z(a+1,a)+P^{(d)}\\
P_2^{(d)} &\approx \frac{1}{a+n}\left(\frac{1}{n}I_z(a+2,a)-2I_z(a+1,a)+nP^{(d)}\right)\\
P_3^{(d)} &\approx \frac{an(2n-1)}{(2a+2n+1)(a+n)}P^{(d)}+\left(\frac{2a+1}{(2a+2n+1)(a+n)}\right)\left(I_z(a+2,a)-2I_z(a+1,a)\right).
\end{aligned} \tag{S72}$$

These expressions allow us to compute the metrics for a much larger range of $n$ and $d$ values.

#### S2.6 Polygenic score bias for recent genic selection

We provide evidence for the claim made in Section 4 that, "*bias*$_\ell(\tau)$ will reach an equilibrium value that depends approximately on the asymmetry of the detection thresholds at the present day, which in turn, depends on both the timing and strength of selection". We treat the simplest case of a detection threshold $d = 1$, i.e. $\hat{\beta} = \beta$ if the locus is variant in the GWA study sample. The time-varying distribution of the allele frequency is $f_t(\cdot)$, and necessarily depends on the timing and strength of selection. For $t > \tau_s$, the time of the onset of selection, $f_t(z) \propto z^{a-1}(1-z)^{a-1}$. For $t \leq \tau_s$, $f_t(z)$ will be skewed toward one and proportional to $\propto e^{\sigma z} z^{a-1}(1-z)^{a-1}$, where $\sigma = 4Ns$ is the population-scaled selection coefficient. For a larger $\tau_s$, $f_t(z)$ for $t \leq \tau_s$ will have more time to shift toward the stationary distribution under selection.

From Equation (11), we have that $bias_\ell(\tau)$, omitting the locus subscript is,

$$\begin{aligned} bias(\tau) &= \mathbb{E}\left[(\bar{X} - X(\tau))(\beta - \hat{\beta})\right] \\ &= \beta\mathbb{E}\left[(\bar{X} - X(\tau))|\hat{\beta} = 0\right]\mathbb{P}\{\hat{\beta} = 0\} \\ &= \beta\left(\mathbb{E}\left[\bar{X}|\hat{\beta} = 0\right] - \mathbb{E}\left[X(\tau)|\hat{\beta} = 0\right]\right)\mathbb{P}\{\hat{\beta} = 0\} \\ &= \beta\left(-\mathbb{P}\{\bar{X} = -1|\hat{\beta} = 0\} + \mathbb{P}\{\bar{X} = +1|\hat{\beta} = 0\} - \mathbb{E}\left[X(\tau)|\hat{\beta} = 0\right]\right)\mathbb{P}\{\hat{\beta} = 0\}. \end{aligned} \tag{S73}$$

For large $\tau$, $\mathbb{E}\left[X(\tau)|\hat{\beta} = 0\right] \to 0$, such that,

$$bias(\tau) \to \beta\left(\mathbb{P}\{\bar{X} = +1|\hat{\beta} = 0\} - \mathbb{P}\{\bar{X} = -1|\hat{\beta} = 0\}\right)\mathbb{P}\{\hat{\beta} = 0\}, \tag{S74}$$

which shows that the $bias(\tau)$ will equilibrate at some value that depends on the difference between the $+$ and $-$ detection thresholds as well as the probability that $\hat{\beta} = 0$. This difference, in turn, depends on the time of the onset of selection and the selection coefficient (relative to the mutation rate) itself.

##### S2.7 Necessary moments, under neutrality and at stationarity

We provide analytic expressions for the moments which constitute the various metrics under the assumption of equal mutation rates. In addition, we provide simplified expressions when the detection thresholds are equal.

**Moments of the population allele frequency and genotype.** Because the population is at stationarity, the moments in this subsection are time-invariant. They require integration over the stationary density of the population allele frequency, which is beta-distributed, and in (b) require integration over the Hardy-Weinberg sampling process.

a. *Moments of the population allele frequency:*

$\mathbb{E}\left[Z_\ell\right] = \frac{1}{2}$, $\mathbb{E}\left[Z_\ell^2\right] = \frac{a+1}{2(2a+1)}$, and $\mathbb{E}\left[Z_\ell(1 - Z_\ell)\right] = \frac{a}{2(2a+1)}$.

b. *Moments of a genotype:*

$\mathbb{E}\left[X_{i\ell}\right] = 0$ and thus $\mathbb{V}\left[X_{i\ell}(t)\right] = \mathbb{E}\left[X_{i\ell}^2(t)\right] = \frac{a+1}{2a+1}$.

**Moments specific to the GWA study.** These moments require integration over the stationary density of the population allele frequency and the sampling probabilities for a sample of $n$ individuals.

a. *Moments of the mean genotype in the GWA study sample:*

$\mathbb{E}\left[\bar{X}_\ell\right] = 0$ and $\mathbb{E}\left[\bar{X}_\ell^2\right] = \frac{1}{n}\left(\frac{a+n}{2a+1}\right)$.

b. *Product of the mean genotype in the GWA study sample and the effect estimate:*

$$\begin{aligned}\mathbb{E}\left[\bar{X}_\ell\hat{\beta}_\ell\right] &= \mathbb{E}\left[\mathbb{E}\left[\bar{X}_\ell\hat{\beta}_\ell|\bar{X}_\ell\right]\right] = \beta_\ell\mathbb{E}\left[\bar{X}_\ell\mathbb{1}_{\{\bar{X}_\ell\in(\gamma-1,1-\gamma)\}}\right] \\ &= \beta_\ell\sum_{i=d_{\ell1}}^{2n-d_{\ell2}}\left(\frac{i-n}{n}\right)\binom{2n}{i}\mathbb{E}\left[Z_\ell^i\left(1-Z_\ell\right)^{2n-i}\right] \\ &= \beta_\ell\sum_{i=d_{\ell1}}^{2n-d_{\ell2}}\left(\frac{i-n}{n}\right)\binom{2n}{i}\frac{B(a+i,b+2n-i)}{B(a,b)}.\end{aligned} \tag{S75}$$

And, for $d_{\ell1}=d_{\ell2}$,

$$\mathbb{E}\left[\bar{X}_\ell\hat{\beta}_\ell\right] = 0. \tag{S76}$$

And,

$$\mathbb{E}\left[\bar{X}_\ell^2\hat{\beta}_\ell\right] = 2\beta_\ell\sum_{i=d_{\ell1}}^{n-d_{\ell2}}\left(\frac{i-n}{n}\right)^2\binom{2n}{i}\frac{B(a+i,a+2n-i)}{B(a,a)} \tag{S77}$$

For $d_{\ell1}=d_{\ell2}=d_\ell$,

$$\begin{aligned}\mathbb{E}\left[\bar{X}_\ell^2\hat{\beta}_\ell\right] &= \beta_\ell\left(\frac{a+n}{n(2a+1)} - 2\sum_{i=0}^{d_\ell-1}\left(\frac{i-n}{n}\right)^2\binom{2n}{i}\frac{B(a+i,a+2n-i)}{B(a,a)}\right) \\ &= \beta_\ell\left(\frac{a+n}{n(2a+1)} - 2P_1^{(d_\ell)}\right).\end{aligned} \tag{S78}$$

Under our simple threshold model, the corresponding second moment of $\hat{\beta}_\ell$ is equal to the previous expression multiplied by $\beta_\ell$.

c. *First moment of the mean phenotype in the GWA study sample:* $\mathbb{E}\left[\bar{Y}\right]=0$.

d. *First moment of the estimated intercept term:*

$$\mathbb{E}\left[\hat{C}\right] = \mathbb{E}\left[\bar{Y}\right] - \sum_{\ell=1}^{L}\mathbb{E}\left[\bar{X}_\ell\hat{\beta}_\ell\right] = C - \sum_{\ell=1}^{L}\beta_\ell\sum_{i=d_{\ell1}}^{2n-d_{\ell2}}\left(\frac{i-n}{n}\right)\binom{2n}{i}\frac{B(a+i,a+2n-i)}{B(a,a)}, \tag{S79}$$

which, for equal detection thresholds equals 0.

**Moments involving both the ancient and contemporary genotypes.** These moments involve quantities from two time points: the ancient sampling time $\tau$ and the GWA study at the present. To compute these moments, we use the spectral representation of the *tdf* (Supplementary Text S1.6).

a. *Product of the first moments of the ancient and contemporary mean genotype:*

$$\begin{aligned}\mathbb{E}\left[X_\ell(\tau)\bar{X}_\ell(0)\right] &= \frac{1}{n}\sum_{i=1}^{n}\mathbb{E}\left[\mathbb{E}\left[X_\ell(\tau)X_{i\ell}(0)|Z_\ell(0),Z_\ell(\tau)\right]\right] \\ &= \mathbb{E}\left[(2Z_\ell-1)(2Z_\ell(\tau)-1)\right] \\ &= \frac{1}{B(a,b)}\sum_{k=0}^{1}\frac{e^{-\lambda_k\tau}}{\langle B_k,B_k\rangle_\pi}\langle 2z-1,B_k\rangle_\pi^2 \\ &= e^{-a\tau}\left(\frac{1}{2a+1}\right).\end{aligned} \tag{S80}$$

b. *Product of the moments of the ancient and contemporary mean genotypes, and the effect estimate:*

$$\begin{aligned}\mathbb{E}\left[\bar{X}_\ell\hat{\beta}_\ell X_\ell(\tau)\right] &= \sum_{i=d_{\ell 1}}^{2n-d_{\ell 2}} \left(\frac{i-n}{n}\right)\binom{2n}{i}\sum_{k=0}^{1}\frac{e^{-\lambda_k\tau}}{\langle B_k, B_k\rangle_\pi}\langle 2z-1, B_k\rangle_\pi\langle z^i(1-z)^{2n-i}, B_k\rangle_\pi \\ &= e^{-a\tau}\beta_\ell\sum_{i=d_\ell}^{2n-d_\ell}\left(\frac{(i-n)^2}{n(a+n)}\right)\binom{2n}{i}\frac{B(a+i,a+2n-i)}{B(a,a)}.\end{aligned} \tag{S81}$$

For equal detection thresholds, $d_\ell$, and using the variables defined in Equation (S35), we have,

$$\mathbb{E}\left[\bar{X}_\ell\hat{\beta}_\ell X_\ell(\tau)\right] = \beta_\ell e^{-a\tau}\left(\frac{1}{2a+1} - 2P_2^{(d_\ell)}\right). \tag{S82}$$

The corresponding second moment of $\hat{\beta}_\ell$ is equal to the previous expression multiplied by $\beta_\ell$.

**Moments involving the ancient (but not contemporary) genotype**. These moments involve the ancient genotype $X_\ell(\tau)$ and the contemporary effect estimate $\hat{\beta}_\ell$, but not the contemporary genotypes.
a. *Product of the first moments of the ancient genotype and the effect estimate:*

$$\begin{aligned}\mathbb{E}\left[X_\ell(\tau)\hat{\beta}_\ell\right] &= \frac{1}{B(a,a)}\sum_{i=d_{\ell 1}}^{2n-d_{\ell 2}}\binom{2n}{i}\sum_{k=0}^{1}\frac{e^{-\lambda_k\tau}}{\langle B_k, B_k\rangle_\pi}\langle z^i(1-z)^{2n-i}, B_k\rangle_\pi\langle 2z-1, B_k\rangle_\pi \\ &= e^{-a\tau}\sum_{i=d_{\ell 1}}^{2n-d_{\ell 2}}\binom{2n}{i}\left(\frac{B(a+i,a+2n-i)}{B(a,a)}\right)\left(\frac{i-n}{a+n}\right) = 0,\end{aligned} \tag{S83}$$

for equal thresholds $d_{\ell 1} = d_{\ell 2}$. This result is due to the fact that, in Equation (S83), the $i$-th term is equal to the $(2n-i)$-th term (and the $n$-th term is 0).
b. *Product of the second moments of the ancient genotype and first moment of the effect estimate:*

$$\begin{aligned}\mathbb{E}\left[X_\ell^2(\tau)\hat{\beta}_\ell\right] &= \frac{\beta_\ell}{B(a,b)}\sum_{i=d}^{2n-d}\binom{2n}{i}\sum_{k=0}^{2}\frac{e^{-\lambda_k\tau}}{\langle B_k, B_k\rangle_\pi}\langle 1-2z+2z^2, B_k\rangle_\pi\langle z^i(1-z)^{2n-i}, B_k\rangle_\pi \\ &= \frac{\beta_\ell}{2a+1}\sum_{i=d}^{2n-d}\binom{2n}{i}\frac{B(a+i,a+2n-i)}{B(a,a)} \\ &\quad\times\left[(a+1) + e^{-(2a+1)\tau}\left(\frac{(2a+1)i(i-2n)+an(2n-1)}{(2a+2n+1)(a+n)}\right)\right].\end{aligned} \tag{S84}$$

For equal detection thresholds $d_\ell$, and Using the terms defined in Equation (S35), we can express Equation (S84) more succinctly,

$$\mathbb{E}\left[X_\ell^2(\tau)\hat{\beta}_\ell\right] = \frac{\beta_\ell}{2a+1}\left[(a+1)(1-2P^{(d_\ell)}) - 2e^{-(2a+1)\tau}P_3^{(d_\ell)}\right]. \tag{S85}$$

The moment $\mathbb{E}\left[X_\ell^2(\tau)\hat{\beta}_\ell^2\right]$ is equal to the previous expression multiplied by $\beta_\ell$.

### References

1. Simons YB, Bullaughey K, Hudson RR, Sella G. A population genetic interpretation of GWAS findings for human quantitative traits. PLoS Biology. 2018;16. https://doi.org/10.1371/journal.pbio.2002985.

2. Sham PC, Purcell SM. Statistical power and significance testing in large-scale genetic studies. Nature Reviews Genetics. 2014;15(5):335–346. https://doi.org/10.1038/nrg3706.

3. Wang Y, Guo J, Ni G, Yang J, Visscher PM, Yengo L. Theoretical and empirical quantification of the accuracy of polygenic scores in ancestry divergent populations. Nature Communications. 2020;11(1). https://doi.org/10.1038/s41467-020-17719-y.

4. Griffiths RC, Spano D. Diffusion processes and coalescent trees. arXiv. 2010. http://arxiv.org/abs/1003.4650.

5. Song YS, Steinrücken M. A simple method for finding explicit analytic transition densities of diffusion processes with general diploid selection. Genetics. 2012;190(3):1117–1129. https://doi.org/10.1534/genetics.111.136929.
